# Multiomics responses to seasonal variations in diel cycles in the marine phytoplanktonic picoeukaryote *Ostreococcus tauri*

**DOI:** 10.1101/2023.07.31.551326

**Authors:** Ana B. Romero-Losada, Christina Arvanitidou, M. Elena García-Gómez, María Morales-Pineda, M. José Castro-Pérez, Mercedes García-González, Francisco J. Romero-Campero

## Abstract

Earth tilted rotation and translation around the Sun produce one of the most pervasive periodic environmental signals on our planet giving rise to seasonal variations in diel cycles. Although marine phytoplankton plays a key role on ecosystems and present promising biotechnological applications, multiomics integrative analysis of their response to these rhythms remains largely unexplored. We have chosen the marine picoeukaryote *Ostreococcus tauri* as model organism grown under summer long days, winter short days, constant light and constant dark conditions to characterize these responses in marine phytoplankton. Although 80% of the transcriptome present diel rhythmicity under both seasonal conditions less than 5% maintained oscillations under all constant conditions. A drastic reduction in protein abundance rhythmicity was observed with 55% of the proteome oscillating. Seasonally specific rhythms were found in key physiological processes such as cell cycle progression, photosynthetic efficiency, carotenoid content, starch accumulation and nitrogen assimilation. A global orchestration between transcriptome, proteome and physiological dynamics was observed with specific seasonal temporal offsets between transcript, protein and physiological peaks.

## Introduction

Marine phytoplankton plays a pivotal role in Earth’s ecosystems, acting as primary producers by contributing to approximately 45% of global photosynthetic net primary production^1, 2^. Consequently, they not only sustain the existence of most oceanic life but also support life across the entire planet. Furthermore, throughout the evolutionary history of life on Earth, oceanic phytoplankton have exerted a profound influence^3^. For instance, the emergence of eukaryotic phytoplankton during the Cryogenian period, characterized by near complete glaciation, possibly played a critical role in the rise of oceanic oxygen and nutrient levels. This may have subsequently led to the Cambrian explosion, a fundamental evolutionary event in the animal kingdom^4, 5^.

Marine phytoplankton’s growth, development and biomass production are coupled with CO_2_ fixation, making them promising sustainable sources for various biotechnological applications, particularly relevant in the context of the current climatic crisis. These applications encompass biostimulants in agriculture, biodiesel production, wastewater and contaminants bioremediation, cosmetics and nutraceuticals^6–8^. However, despite their potential, many of these applications remain economically unfeasible at the industrial level^9,10^. Hence, optimizing culture systems and production methods is imperative, necessitating the application of artificial intelligence and machine learning tools^11^, along with further characterization of the molecular mechanisms underpinning microalgae physiology using systems and synthetic biology techniques^12–15^.

Light availability is a critical environmental condition with a profound impact on marine phytoplankton growth and physiology^16^. Earth’s rotation produces the most pervasive periodic and rhythmic environmental signal, affecting light on our planet and giving rise to alternating cycles of day (light periods or photoperiods) and night (dark periods or skotoperiods), collectively known as diel cycles. Earth’s tilted rotational axis and its translation around the Sun also lead to seasonal variations in diel cycles resulting in long days (long photoperiods) with short nights (short skotoperiods) in summer, and short days (short photoperiods) with long nights (long skotoperiods) in winter. Seasonality has been found to play a central regulatory role in marine phytoplankton dynamics^17–20^. However, the molecular mechanisms underpinning these responses are not yet characterized.

Chronobiology is an emerging multidisciplinary field in biology that focuses on the study of the timing and regulation of biological rhythms and their synchronization with the previously described 24 hour diel cycles, known as circadian rhythms^21^. The molecular mechanisms that have evolved to anticipate and respond to these rhythms are referred to as circadian clocks. They are constituted by autonomous oscillating molecular systems that can be entrained by external cyclic inputs, such as diel cycles, and produce rhythmic output of biological processes. The distinctive characteristic of *bona fide* circadian biological rhythms is that they are self-sustained maintaining rhythmicity under constant conditions. To identify these biological rhythms, chronobiology experiments are typically designed as a sequence of several consecutive days under alternating light/dark cycles, followed by several consecutive days when light or dark is kept constant, called free-running conditions^21^. While extensive chronobiological studies on circadian rhythms at the molecular and physiological levels have been conducted in many model organisms^22–31^, only initial steps have been taken to analyze such systems in marine phytoplankton using omics technologies^32–34^.

To characterize the responses of marine phytoplankton to seasonal variations in diel cycles, we selected the model phytoplanktonic picoeukaryote *Ostreococcus tauri* (Ostreococcus) due to its cellular simplicity^35–37^,availability of its fully sequenced and annotated genome^38–41^, its position in the evolutionary history of the green lineage^42–45^ and its abundance in marine ecosystems^18, 46–48^. Intensive and extensive studies on the physiology of Ostreococcus have provided a solid foundation for systems biology and omics studies. These include circadian chronobiological studies^49–52^, cell cycle analysis^53–56^, sexual reproduction^57^, viral infections dynamics^58, 59^, biomass composition analysis^60–65^ and omics analysis^58, 66–70^. Nevertheless, Ostreococcus responses to seasonal variations in diel cycles, as well as free running conditions, remain to be explored using integrative multiomics analysis.

In this study, in order to characterize Ostreococcus transcriptome, proteome and physiological rhythmicity we cultivated Ostreococcus using photochemostats operated in continuous regime, simulating summer long day conditions (LD, 16h light : 8h dark) and winter short day conditions (SD, 8h light : 16h dark), (Supplementary Fig. 1). Photoperiod light intensity fluctuations were gradual, mimicking diurnal cycles in light variations. We collected samples for three consecutive days every four hours (ZT0, ZT4, ZT8, ZT12, ZT16 and ZT20) where ZTN (Zeitgeber time N) marks the time point N hours after the beginning of the light period. Transcriptomic, proteomic, and physiological data were generated from these samples. Subsequently, to determine *bona fide* circadian genes, we transferred cultures to free-running conditions consisting of constant light (LL) and dark (DD). No samples were collected during the first day to allow cultures acclimation, and then we collected samples every four hours for two consecutive days to generate transcriptomic data. In this study, by integrating transcriptomic, proteomic data with physiological measurements, we unveiled a seasonal adaptive global orchestration between transcriptome, proteome, and physiological dynamics, with specific seasonal temporal offsets between transcript, protein and physiological peaks.

## Results

### Transcriptome rhythmicity under different seasonal variations of diel cycles and identification of *bona fide* circadian genes

Robust global rhythmicity was detected in the transcriptomes of our cultures by using Hierarchical Clustering (HC) over our RNA-seq data corresponding to the 36 time points collected under LD and SD conditions. Three distinct clusters were identified corresponding to midday (LD ZT4, SD ZT4 and LD ZT8), dusk (SD ZT8, LD ZT16 and LD ZT12) and night/dawn with two clear subclusters distinguishing between LD and SD (LD ZT0 and LD ZT20 on one hand and SD ZT12, SD ZT16, SD ZT20 and ZT0 on the other hand). This reveals a clear distinction between the night/dawn LD and SD transcriptomes. Accordingly, Principal Components Analysis (PCA) revealed a cyclic circular organization of the transcriptomes over diel cycles, being more apparent under LD than SD conditions (Supplementary Fig. 1). This suggests more complex gene expression patterns under SD than LD. The non-parametric methods for rhythmicity implemented in the Bioconductor R package RAIN^71^ were used to identify rhythmic genes under the different conditions of interest in this study (Figure 1A). Seasonal variations in photoperiod did not affect transcriptome rhythmicity, as the sets of rhythmic genes under LD and SD entrainment were almost coincident, comprising approximately 80% of the genome under both conditions. This nearly complete entire rhythmic transcriptome is in agreement with previous results obtained under neutral day conditions (ND, 12h light / 12 hours dark) for *Ostreococcus* using microarray data^51^ and other chlorophyte microalgae like *Chlamydomonas reinhardtii* using RNA-seq data^72^. The non rhythmic genes in our experiments were either completely repressed or very lowly expressed. Indeed, rhythmic genes presented significant maximum expression levels three times greater than non-rhythmic genes according to a p-value of 1.45×10^-4^ computed using Mann-Whitney-Wilcoxon test. They were mainly involved in stress responses such as viral infection^58^. These genes could also be rhythmic once the corresponding signal activates them which would result in a fully rhythmic transcriptome under alternating light/dark cycles.

**Figure 1.**
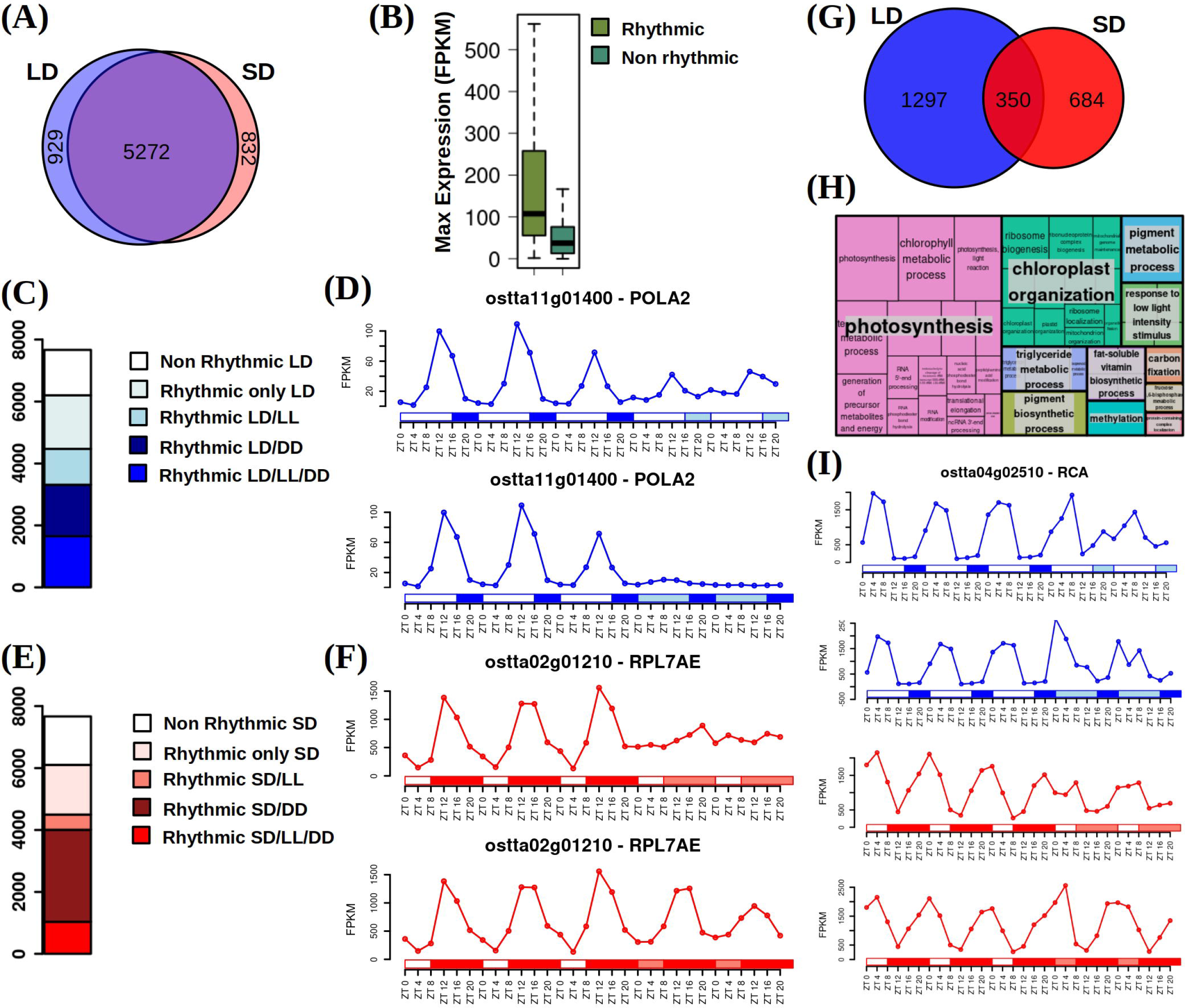
Transcriptome rhythmicity under alternating light/dark cycles and identification of *bona fide* circadian genes including free running conditions. **(A)** Venn diagram comparing rhythmic genes under long day (LD, 16h light / 8h dark) conditions (light blue circle) and short day (SD, 8h light / 16h dark) conditions (light red circle). Genes exhibiting rhythmic expression patterns are almost identical under both photoperiodic entrainments. **(B)** Boxplot representing the maximum expression level of rhythmic genes (light green) and non-rhythmic genes (dark green). Medians are represented by central horizontal lines, upper and lower quartiles by boxes, minimum and maximum values by whisker ends. Rhythmic genes present significant maximum expression levels three times greater than non-rhythmic genes according to a p-value of 1.45 × 10^-4^ computed using Mann-Whitney-Wilcoxon test. Gene expression levels are measured as FPKM (Fragments Per Kilobase of transcript per Million fragments mapped). **(C)** Barplot representing with blue colors different rhythmic gene sets under LD conditions. From bottom to top: circadian genes exhibiting rhythmicity under LD, constant light (LL) and constant dark (DD); rhythmic genes under LD and DD requiring a dark input; rhythmic genes under LD and LL requiring a light input; rhythmic genes only under LD requiring the alternation of light/dark periods and non-rythmic genes. **(D)** Gene expression profiles under LD, LL and DD of *DNA polymerase alpha subunit B* (*ostta11g01400*, *POLA2*). White rectangles represent photoperiods (light periods or days), blue filled rectangles correspond to skotoperiods under LD (dark periods or nights), light blue rectangles mark subjective days or nights under LL and DD respectively after LD entrainment. ZTN, Zeitgeber time N, marks the time point N hours after dawn (lights on, ZT0). *POLA2* illustrates that specific genes involved in DNA replication require light maintaining rhythmicity under LL whereas being strongly repressed under DD. **(E)** Barplot representing with red colors different rhythmic gene sets under SD conditions. From bottom to top: circadian genes exhibiting rhythmicity under SD, LL and DD; rhythmic genes under SD and DD requiring a dark input; rhythmic genes under SD and LL requiring a light input; rhythmic genes only under SD requiring the alternation of light/dark periods and non-rhythmic genes. **(F)** Gene expression profiles under SD, LL and DD of *Ribosomal protein L7Ae* (*ostta02g01210*, *RPL7AE*). White rectangles represent photoperiods (light periods or days), red filled rectangles correspond to skotoperiods under SD (dark periods or nights), light red rectangles mark subjective days or nights under LL and DD respectively after SD entrainment. *RPL7AE* illustrates that specific genes involved in translation require dark maintaining rhythmicity under DD whereas presenting flat expression levels under LL. **(G)** Venn diagram comparing circadian genes identified after LD entrainment (blue circle) and after SD entrainment (red circle). Only a reduced number of genes are identified as *bona fide* circadian maintaining rhythmicity under the free running conditions LL and DD after both LD and SD entrainments. **(H)** Treemap summarizing the biological processes significantly enriched over the *bona fide* circadian genes or rhythmic genes under LD, SD, LL and DD. Rectangle sizes represent significance levels. Semantically similar biological processes are grouped together into the same colored rectangles. The most representative biological process is shown for each rectangle. **(I)** Gene expression profiles under LD, SD, LL and DD of *RuBisCO Activase* (*ostta04g02510*, *RCA*) exemplifying that *bona fide* circadian genes that maintain rhythmicity under light/dark cycles as well as free running conditions are involved in photosynthesis related processes.

After entrainment under LD and SD conditions, in order to distinguish between circadian genes and those responding to the rhythmic changes between light periods (photoperiods) and dark periods (skotoperiods), our cultures were transferred to free-running conditions consisting of constant light (LL) and constant dark (DD). A clear reduction in rhythmicity, dependent on the previous entrainment, was observed with LL having a more detrimental effect than DD (Figure 1C,E). Approximately 21% of the transcriptome was completely reliant on the alternation of photoperiods and skotoperiods to maintain rhythmicity since these genes lost their rhythms under both LL and DD independently from the entrainment regime. In contrast, the transcriptome proportion that maintained its oscillations exclusively under LL or DD was dependent on the previous entrainment. Whereas 36.6% kept cycling under LL after LD entrainment, only 20% were rhythmic under LL after SD entrainment. The oscillating genes under LL, both after LD and SD entrainment, were found to be significantly involved in DNA replication and Chromosome organization (Supplementary Fig. 2). Key genes implicated in DNA replication such as *DNA polymerase alpha subunit B* (*ostta11g01400*, *POLA2), Minichromosome Maintenance 6* (*MCM6*, *ostta01g02580*) and *Proliferating Cell Nuclear Antigen* (*PCNA*, *ostta06g02890*) maintained oscillations under LL but were strongly repressed under DD (Figure 1D, Supplementary Fig. 2). The detrimental effect of DD over transcriptome rhythmicity was smaller than that of LL, with almost 43.2% maintaining oscillations after LD entrainment and a remarkable 52.2% after SD entrainment. The biological processes RNA processing and ribosome biogenesis were found to be significantly enriched among the rhythmic genes under DD, both after LD and SD entrainment (Supplementary Fig. 3). Important genes in ribosome assembly such as *Ribosomal protein L7Ae* (*ostta02g01210*, *RPL7AE*), *U3 small nucleolar RNA-associated proteins 11* and *14 (Utp11, ostta06g01560*; *Utp14*, *ostta04g00770*) presented clear periodic oscillations under DD whereas under LL a flat non-zero profile was observed (Figure 1F, Supplementary Fig. 3).

We consider *bona fide* circadian genes those presenting rhythmicity under both LD and SD as well as maintaining their rhythmic expression profiles under both LL and DD free-running conditions. A clear dependence on the previous entrainment regime was observed with 1647 and 1034 genes keeping rhythmicity under both free running conditions after LD and SD entrainment respectively. These two sets overlapped partially identifying 350 *bona fide* circadian genes corresponding to 4.56% of the *Ostreococcus* transcriptome (Figure 1G). Functional enrichment analysis revealed that these genes were significantly involved in photosynthesis, chloroplast organization and pigment metabolic process (Figure 1H). For example, the gene codifying for *RuBisCO Activase* (*ostta04g02510*, *RCA*) was observed to present rhythmic expression profiles under LD, SD, LL and DD. This gene has been reported to be regulated by the circadian clock in many diverse plant species such as Arabidopsis, tomato, tobacco, apple, rice and wheat although no data under free running conditions were analysed. In the case for *Ostreococcus* our results indicate that this gene is *bona fide* circadian being predominantly regulated by the endogenous autonomous system constituting the circadian clock in this species since it maintains rhythmicity independently from the external signals.

### Effects of free-running conditions over rhythmic gene expression patterns

These rhythmic gene sets were consistent with the ones obtained using the co-sinusoidal parameterized methods implemented in the R package circacompare^73^ showing the robustness of our results, Supplementary Fig. 4. These models represent, for each rhythmic gene expression profile, its phase (the time point when the maximum expression level is reached) and amplitude (the range of gene expression from the minimum to the maximum value). Moreover, these methods allow us to assess the statistical significance of the differences in phase and amplitude between different expression profiles. We used this approach to identify the effects of free-running conditions, LL and DD, over the rhythmic gene expression patterns emerged under LD and SD. In the LD entrained cultures we detected significant reductions in amplitude when transferred to both LL and DD being more drastic under LL according to p-values of 4.23×10^-140^ and 1.09 × 10^-60^ respectively (Figure 2A). In SD entrained cultures amplitude reduction was only noted when transferred to LL with a p-value of 5.94×10^-65^. Nonetheless a slight but significant amplitude increase was found when transfer to DD with p-value of 2.5× 10^-8^. These changes in amplitude could possibly be due to culture synchrony disruption at the transcriptomics level. Specifically, amplitude reductions could be associated with the transcriptional programs of individual cells becoming desynchronized such as individual gene expression profiles become largely out of phase. This would result in damped average gene expression oscillations at the level of the entire cell culture (Supplementary Fig. 4). Individual cell desynchronization resulting in damped oscillations have been reported previously in Arabidopsis leaves grown under constant light^74, 75^. In contrast, a synchrony increase between individual cell transcriptional programs would make individual gene expression profiles get in phase producing an increase in amplitude in the average gene expression oscillations at the entire cell culture (Supplementary Fig. 4). Therefore our data indicate that in Ostreococcus cultures LL has a greater desynchronizing effect than DD suggesting the need for a dark period to maintain culture synchrony. Even a synchrony increase in SD entrained cultures was observed when they were transferred to DD free-running conditions.

**Figure 2.**
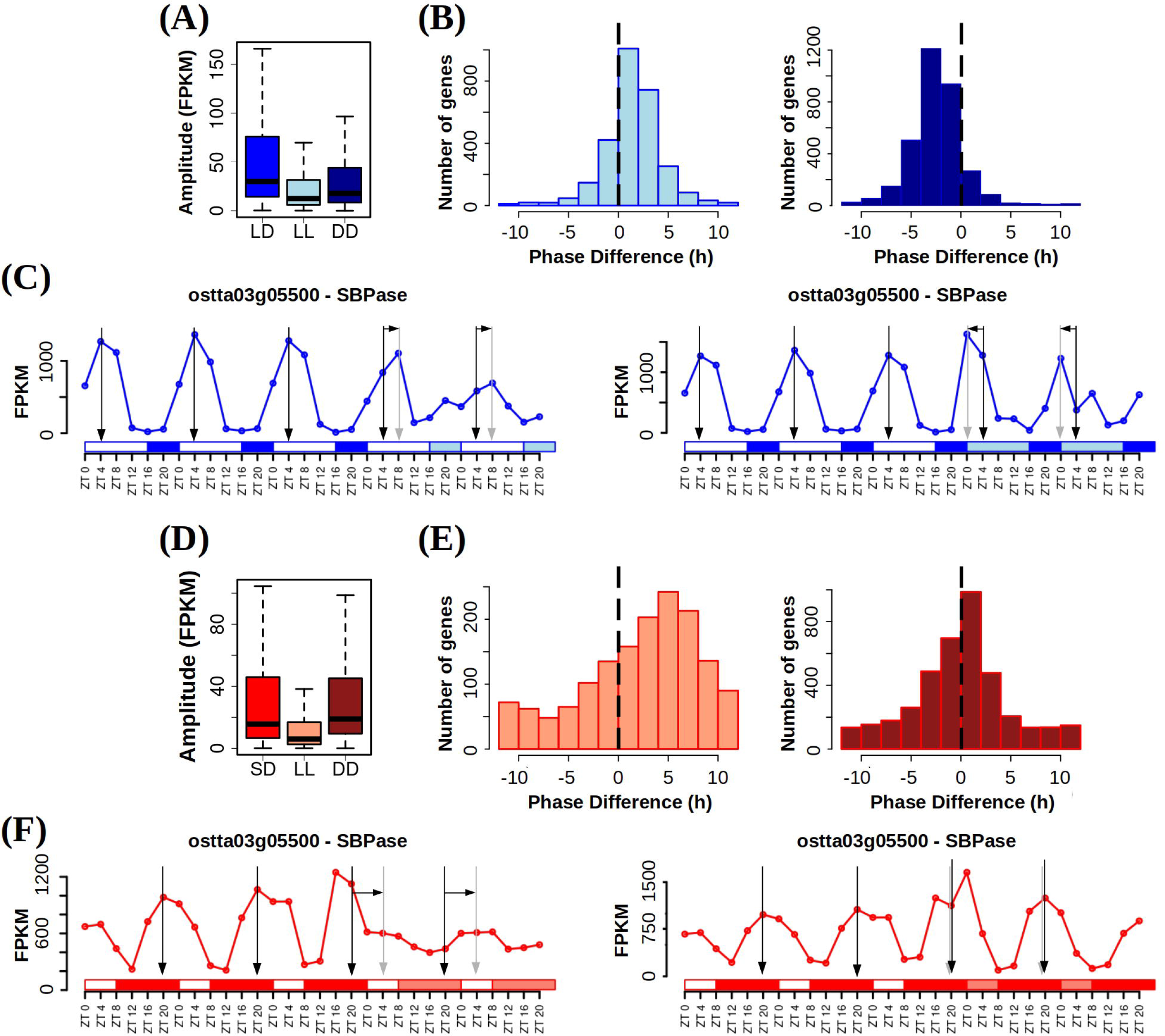
Free running conditions effects over gene expression profiles. **(A)** Boxplot representing rhythmic genes amplitude or maximum expression level reached under long day conditions (LD, alternating 16h light / 8h dark, blue), when cultures were transferred to free running conditions consisting of constant light (LL, light blue) and when cultures were transferred to free running conditions consisting of constant dark (DD, dark blue) after LD entrainment. Medians are represented by central horizontal lines, upper and lower quartiles by boxes, minimum and maximum values by whisker ends. LL and DD amplitudes are significantly reduced with respect to LD according to p-values of 4.23 × 10^-140^ and 1.09 × 10^-60^ possibly due to a decline in culture synchrony under free running conditions. LL amplitudes are also reduced when compared to DD according to a p-value of 6.71 × 10^-28^ suggesting more severe loss of rhythmicity under LL than DD. P-values were computed using Mann-Whitney-Wilcoxon test. Gene expression levels are measured as FPKM (Fragments Per Kilobase of transcript per Million fragments mapped). **(B)** Histograms showing the distribution of the number of genes exhibiting positive and negative shifts in phase or maximum expression level time point when cultures are transferred from LD to free running conditions consisting in LL (light blue, left) and DD (dark blue, right). Vertical dashed lines mark no shift. Positive forward phase shifts are observed when cultures are transferred from LD to LL whereas negative backward phase shifts are apparent when transferred to DD. **(C)** Gene expression profiles under LD, LL and DD of *Sedoheptulose-bisphosphatase* (*ostta03g05500*, *SBPase*). White rectangles represent photoperiods (light periods or days), blue filled rectangles correspond to skotoperiods under LD (dark periods or nights), light blue rectangles mark subjective days or nights under LL and DD respectively after LD entrainment. ZTN, Zeitgeber time N, marks the time point N hours after dawn (lights on, ZT0). Vertical black arrows mark LD phases, vertical grey arrows mark LL and DD phases and horizontal black arrows represent phase shifts. *SBPase* illustrates how genes after LD entrainment present reduced amplitudes under LL and DD, forward phase shifts under LL and backward phase shift under DD being these changes more drastic under LL than DD. **(D)** Boxplot representing rhythmic genes amplitude or maximum expression level reached under short day conditions (SD, alternating 8h light / 16h dark, red), when cultures were transferred to free running conditions consisting of constant light (LL, light red) and when cultures were transferred to free running conditions consisting of constant dark (DD, dark red) after SD entrainment. LL amplitudes are significantly reduced with respect to SD according to a p-value of 5.94 × 10^-65^ possibly due to a decline in culture synchrony under LL. In contrast, DD amplitudes present a slight significant increase when compared to SD according to a p-value of 2.5 × 10^-8^ suggesting an increase in culture synchrony under DD. P-values were computed using Mann-Whitney-Wilcoxon test. **(E)** Histograms showing the distribution of the number of genes exhibiting positive and negative shifts in phase or maximum expression level time point when cultures are transferred from SD to free running conditions consisting in LL (light red, left) and DD (dark red, right). Vertical dashed lines mark no shift. Large positive forward phase shifts are observed when cultures are transferred from SD to LL whereas no substantial phase shifts are apparent when transferred to DD. **(F)** Gene expression profiles under LD, LL and DD of *Sedoheptulose-bisphosphatase* (*ostta03g05500*, *SBPase*). White rectangles represent photoperiods (light periods or days), red filled rectangles correspond to skotoperiods under SD (dark periods or nights), light red rectangles mark subjective days or nights under LL and DD respectively after SD entrainment. Vertical black arrows mark SD phases, vertical grey arrows mark LL and DD phases and horizontal black arrows represent phase shifts. *SBPase* illustrates how genes after SD entrainment present reduced amplitudes and forward phase shifts only under LL with slight increases under DD.

Free-running conditions also affected rhythmic gene expression patterns phases. Independently from the previous entrainment regime, LD and SD, forward phase shifts or advances were detected when cultures were transferred to LL (Figure 2 B and E left). Whereas backward phase shifts or delays were found when cultures were transferred to DD (Figure 2 B and E right). The phase advances in LL were more drastic for SD than LD entrained cultures although the delays in DD were more evident for LD than SD entrained cultures. These types of phase shifts under free-running conditions have been found in behavioral traits in nocturnal animals^76, 77^. This result points to a predominantly nocturnal character of the Ostreococcus transcriptome. Genes involve in photosynthesis such as Sedoheptulose-bisphosphatase (ostta03g05500, SBPase) illustrate for a specific case the responses describe above globally (Figure 2 C and F).

### Effects of seasonal variations in photoperiod over gene expression profiles

To determine the response of rhythmic gene expression profiles to seasonal variations in photoperiod length, we also applied the co-sinusoidal parameterized methods^73^. Precisely, we identified specific differences in phase and amplitude between LD and SD rhythmic gene expression patterns. We observed a significant reduction in amplitude in SD compared to LD entrained cultures, with a p-value of 1.16×10^-111^ (Figure 3A). As previously discussed, this could indicate a decline in culture transcriptional synchronization in SD compared to LD. With regard to the effects on gene expression phases or time points of maximum gene expression level, under both LD and SD, we observed an accumulation during the dark periods or skotoperiods with most genes peaking during nights (Figure 3B). This finding supports the nocturnal character of the Ostreococcus transcriptome. We also noted an anticipation of SD phases with respect to LD phases. Whereas under LD entrainment, gene phases accumulate uniformly from the end of the day ZT12, to the end of the night ZT20, under SD entrainment gene phases are concentrated during the first half of the night from ZT8 to ZT16. Genes involved in cell cycle progression and metabolism, such as *Cyclin B* (*ostta01g06150*, *CYCB*) and *Delta-9 acyl-lipid desaturase 1* (*ostta01g00790*, *ADS1*), illustrate the amplitude reductions and phase anticipations reported under SD conditions with respect to LD (Figure 3C).

**Figure 3.**
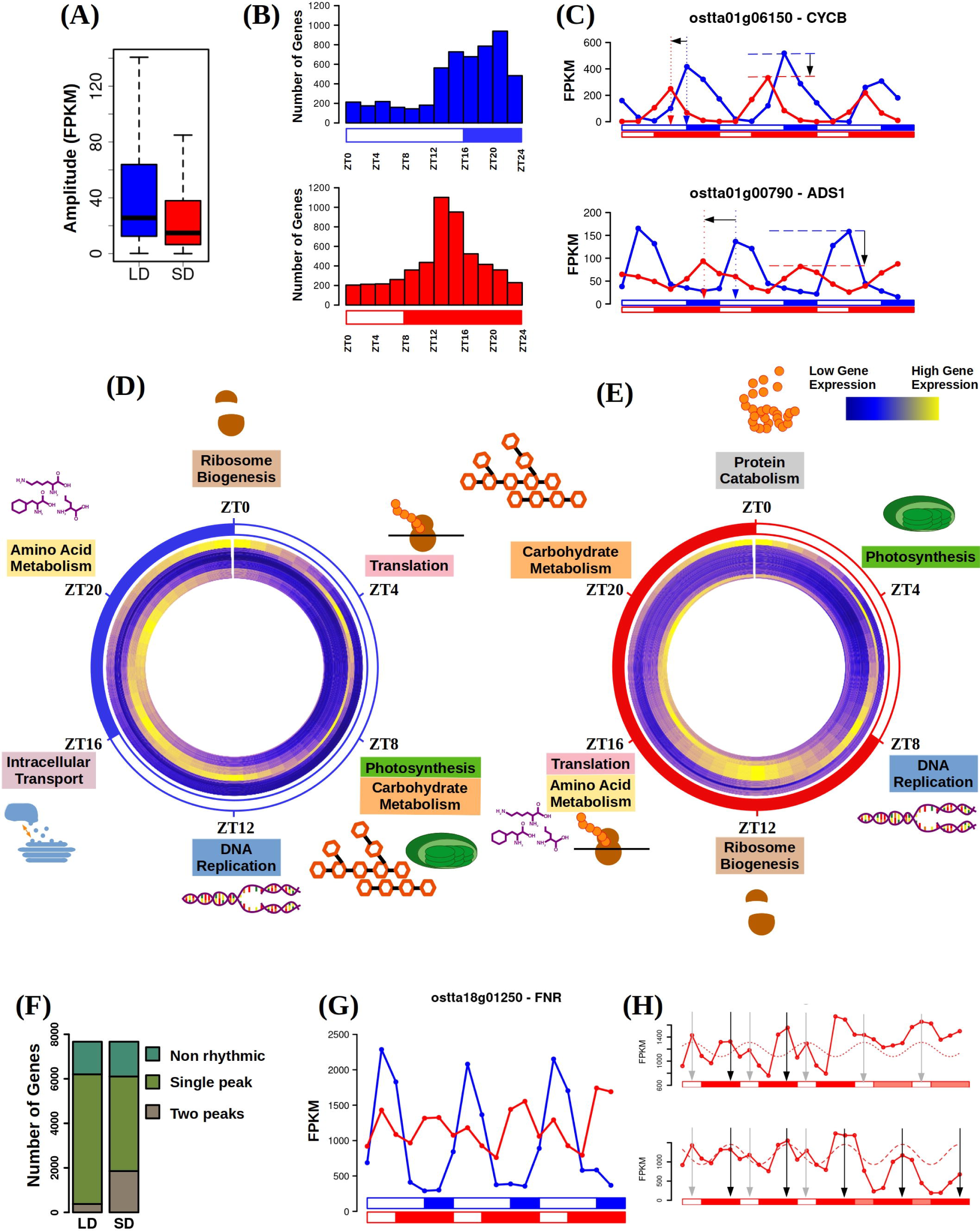
Seasonal and photoperiodic effects over gene expression profiles. **(A)** Boxplot representing rhythmic genes amplitude or maximum expression level reached under long day conditions (LD, alternating 16h light / 8h dark, blue) and under short day conditions (SD, alternating 8h light / 16h dark, red). Medians are represented by central horizontal lines, upper and lower quartiles by boxes, minimum and maximum values by whisker ends. Gene expression levels are measured as FPKM (Fragments Per Kilobase of transcript per Million fragments mapped). SD amplitudes are significantly reduced with respect to LD according to a p-value of 1.16 × 10^-111^ computed using Mann-Whitney-Wilcoxon test possibly due to a decline in culture synchrony under SD. **(B)** Histograms showing the distribution of the number of genes with phase or maximum expression level at specific time points during the day under LD conditions (blue, top) and SD conditions (red, bottom). ZTN, Zeitgeber time N, marks the time point N hours after dawn (lights on). Backward phase shifts are apparent under SD when gene phases are mostly reached around SD midnight ZT12 to ZT16 whereas, under LD, phases are uniformly distributed from dusk (ZT12) to the end of the night. **(C)** Gene expression profiles under LD (blue line) and SD (red line) of *Cyclin B* (*ostta01g06150*, *CYCB*, top) and *Delta-9 acyl-lipid desaturase 1* (*ostta01g00790*, *ADS1*, bottom). White rectangles represent photoperiods (light periods or days), blue and red filled rectangles correspond to skotoperiods under LD and SD respectively (dark periods or nights). Gene expression levels are measured as FPKM (Fragments Per Kilobase of transcript per Million fragments mapped). Blue and red vertical dotted arrows mark LD and SD phases. Horizontal black arrow represents backward phase shifts under SD when compared to LD. Blue and red horizontal dashed lines mark LD and SD amplitudes. Vertical black arrows represent the reductions in amplitude under SD with respect to LD. **(D)** Circular heatmap representing the temporal organization of gene expression profiles under LD conditions. Dark blue stands for low expression whereas yellow represents high expression. Genes are clustered depending on their phase or time of maximum expression being genes with phase at ZT0 located in the outer circle and being genes with phase at subsequent ZTs placed sequentially into inner circles. Biological enriched processes in the gene set with phase at each specific time point are depicting capturing the diurnal transcriptional program over diurnal cycles under LD conditions. **(E)** Similarly, circular heatmap representing the temporal organization of the diurnal transcriptional program under SD conditions. A shift as well as a rearrangement is observed in the SD diurnal transcriptional program when compared to the one inferred for LD. **(F)** Barplot representing in different green colours from top to bottom the number of non rhythmic, single peak rhythmic and two peaks rhythmic genes under LD and SD conditions. An increase in the number of two peaks rhythmic genes is observed as the photoperiod or day shortens (skotoperiod or night lengthens) from LD to SD conditions. **(G)** Gene expression profiles under LD (blue line) and SD (red line) of *Ferredoxin-NADP+ reductase* (*ostta18g01250*, *FNR*). This gene illustrates how specific single peak rhythmic expression patterns under LD conditions become two peaks rhythmic expression profiles under SD conditions. **(H)** Gene expression profiles under SD and free running conditions consisting of constant light (LL) or constant dark (DD) of *Ferredoxin-NADP+ reductase* (*ostta18g01250*, *FNR*). This gene exemplifies how two peaks expression patterns under SD conditions could emerge as the combination of two distinct rhythmic profiles. One depending on the photoperiod (dotted line) with phase marked with a grey vertical arrow maintaining its rhythmicity only under LL (top). Another expression profile is apparent depending on the skotoperiod (dashed line) with phase marked with a black vertical arrow maintaining its rhythmicity only under DD (bottom).

We defined gene clusters based on gene phases or peaking time points. Specifically, for SD and LD entrainment, and for each ZTN (N = 0, 4, 8, 12, 16 or 20), all rhythmic genes under the corresponding conditions reaching their maximum expression level at the given time point were included in a different gene set or cluster (Supplementary Table 3). Functional enrichment analysis was performed on these gene sets to determine significantly enriched biological processes. This identified distinct diel gene activation programs under LD (Figure 3D and Supplementary Fig. 8) and SD (Figure 3E and Supplementary Fig. 9) conditions. These programs consist of different sequences of biological processes activation produced by the previously mentioned gene phase backward shifts under SD with respect to LD. Changes in the activation time point were detected. For example, genes involved in photosynthesis peak at ZT8 under LD and at ZT4 under SD, coinciding with the moment of maximal light irradiance in both cases. Similarly, genes involved in DNA replication reach their maximum expression level at ZT12, end of the day, under LD and at ZT8, the beginning of the night, under SD. We also found rearrangements in the sequence of biological processes activation. For instance, under LD conditions, genes involved in amino acid biosynthesis, ribosome biogenesis, and translation reach their full activation separately at ZT20, ZT0 and ZT4 respectively. However, in SD entrained cultures, first ribosome biogenesis genes are fully activated at ZT12 and subsequently at the same time point, ZT16, amino acid biosynthesis and genes involved in translation reach their maximum expression level.

We developed a co-sinusoidal dynamical model to capture changes in the phase and amplitude of rhythmic gene expression profiles resulting from seasonal changes in photoperiod length. Our model predicts the phase and amplitude of each rhythmic gene for a specific day of the year using a linear interpolation between the phases and amplitudes identified under LD and SD entrainment, Supplementary Fig. 5. To evaluate the predictive power of our model, we used the microarray data generated under neutral day conditions (12 h light : 12 h dark) ^51^. Since gene expression measurements from microarray and RNA-seq data are not directly comparable, only phase prediction could be assessed. Our model successfully predicted the phases of approximately two thirds (63%) of the rhythmic genes with ±4 h error, indicating a linear and gradual adjustment to seasonal changes in photoperiod length in these genes. However, for the remaining one third of the rhythmic genes, a more complex response to photoperiod length is expected.

We observed another response to seasonal variations in photoperiod length related to bimodal rhythmic gene expression profiles with two peaks every day according to apparent 12h periods (Figure 3G). Although two peaks gene profiles were detected in both LD and SD entrained cultures (Supplementary Table 4), a drastic increase in the number of genes with bimodal rhythmicity was observed under SD conditions, 1855 genes, with respect to LD, 376 genes (Figure 3F). Bimodal rhythmicity was not maintained under free running conditions in which distinct single peak profiles were apparent under LL and DD with different phases (Figure 3H). We applied Non Linear Squares (NLS)^78^ to decompose the observed bimodal gene profile in SD entrained cultures into two different single peak profiles one peaking during the photoperiod and another one peaking during the skotoperiod (dark period). We found that under LL free running conditions only the photoperiod peaking profile was maintained whereas under DD conditions only the skotoperiod peaking profile was present. This finding suggests that bimodal rhythmic gene expression emerge from two different regulations, one exerted by the photoperiod and the other one by the skotoperiod. For these bimodal rhythmic genes, we developed a dynamical model that combines two distinct co-sinusoidal profiles to capture changes in the phase and amplitude resulting from seasonal changes in photoperiod length. We observed that under LD conditions these profiles overlap in time producing a single peak profile whereas under SD conditions they become out of phase resulting in a bimodal profile, Supplementary Fig. 6. As a validation, our model predicted the emergence of bimodal rhythmic profiles under neutral conditions, Supplementary Fig. 7.

### Proteome rhythmicity under seasonal variations in photoperiod and integration with transcriptomic rhythmic patterns

Proteins are the primary molecular entities that carry out the majority of biological processes in living cells. Therefore, proteomic analysis, which involves the identification and quantification of proteins, plays a crucial role in characterizing the responses of living organisms to diel cycles^68, 70^. In line with this, we utilized a label-free protein quantification platform that employs independent data acquisition (DIA) called SWATH-MS (sequential windowed acquisition of all theoretical mass spectra)^79^ to generate proteomic data from the same samples used in our transcriptomic analysis conducted under alternating light/dark cycles. Our data successfully identified and quantified 3,672 proteins under LD or SD conditions, accounting for approximately 48% of the *Ostreococcus* annotated proteome, (Supplementary Table 5). The subcellular location of the detected proteins covered most cellular components or organelle. Through data normalization, we were able to eliminate technical biases. Additionally, similar to our transcriptomic data, Principal Components Analysis (PCA) revealed a cyclic circular organization of the proteomes over diel cycles, Supplementary Fig. 10.

The non-parametric methods for rhythmicity implemented in the Bioconductor R package RAIN^71^ were also used to identify proteins with rhythmic abundance profiles under different seasonal variation in photoperiod. A drastic reduction in the number of rhythmic proteins was observed compared to the percentage of rhythmic genes. Specifically, 928 and 1,442 rhythmic proteins were identified under LD and SD conditions, respectively (Supplementary Table 6). Our analysis found that approximately 55% of the detected proteins exhibited rhythmic abundance profiles under either LD or SD, whereas only 9% displayed rhythmicity under both conditions (Figure 4A). The rhythmic protein abundance profiles were compared to the corresponding rhythmic gene expression profiles using the co-sinusoidal parameterized methods implemented in the R package circacompare^73^. Phase offsets between transcript and protein profiles were identified (Supplementary Table 6). Indeed, no global correlation was observed between transcript and protein profiles whereas high global positive correlations were apparent between phase aligned protein and transcript profiles, Supplementary Fig. 11. For instance, the transcript and protein encoded by *Minichromosome Maintenance 2* (*ostta11g00910*, *MCM2*) exhibited a phase offset of eight hours under LD conditions. The maximum gene expression level occurred at ZT12, while the maximum protein abundance was observed at ZT20. Similarly, under SD conditions, there was a phase offset of twelve hours. The maximum gene expression level occurred at ZT8, while the maximum protein abundance was observed at ZT20 (Figure 4B). Specifically, in LD entrained cultures transcript/protein phase offsets presented a unimodal distribution centered around 5 hours, whereas under SD, offsets followed a more uniform distribution, Supplementary Fig. 11. Indeed, the offsets identified under SD were significantly longer than the ones observed in LD according to a p-value of 1.2 × 10 ^-9^ computed using Mann-Whitney-Wilcoxon test. These offsets inferred a diurnal character to the *Ostreococcus* proteome with most proteins reaching their maximum abundance during the day or end of the night, both in LD and SD entrained cultures (Figure 4C). This is in contrast with the nocturnal character exhibited by the transcriptome (Figure 3B).

**Figure 4.**
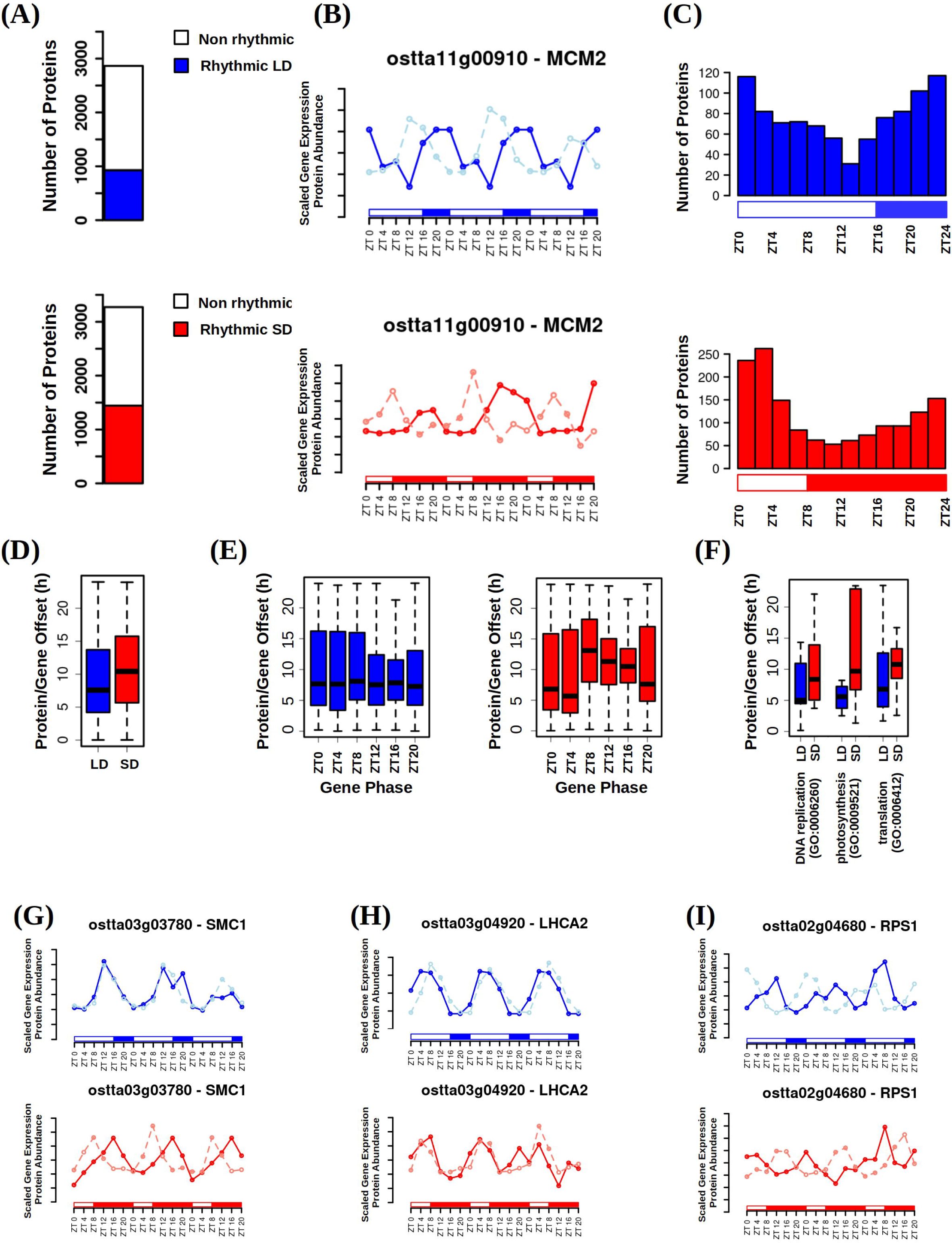
Proteome rhythmicity under alternating light/dark cycles and integration with the transcriptome. **(A)** Barplots representing the number of identified proteins under long day conditions (top, LD, 16h light / 8h dark) and under short day conditions (bottom, SD, 16h light / 8h dark). The number of rhythmic proteins under LD conditions is represented in blue and under SD conditions in red. Non rhythmic proteins are represented in white. **(B)** Protein abundance profiles under LD (top, blue) and SD (bottom, red) conditions represented together with gene expression profiles under LD (top, light blue) and SD (bottom, light red) conditions for *Minichromose Maintenance 2* (*ostta11g00910*, *MCM2*). White rectangles represent photoperiods (light periods or days), blue and red filled rectangles correspond to skotoperiods under LD and SD respectively (dark periods or nights). ZTN, Zeitgeber time N, marks the time point N hours after dawn (lights on, ZT0). *MCM2* illustrates that commonly protein rhythmic profiles exhibit an offset with respect to gene expression profiles. **(C)** Histograms showing the distribution of the number of proteins with phase or maximum abundance at specific time points under LD conditions (top, blue) and SD conditions (bottom, red). Offsets are apparent in protein abundance phases with respect to gene expression phases (time points of maximum protein abundance or gene expression). Under both LD and SD conditions protein abundance phases accumulate at the end of the skotoperiods (dark periods) and during photoperiods (light periods). **(D)** Boxplot representing the offset in hours between protein abundance and gene expression phases under LD (blue) and SD (red) conditions. Medians are represented by central horizontal lines, upper and lower quartiles by boxes, minimum and maximum values by whisker ends. Protein/gene offsets are significantly longer under SD conditions with respect to LD conditions according to a p-value of 1.2 × 10^-9^ computed using Mann-Whitney-Wilcoxon test. **(E)** Boxplots representing protein/gene offsets under LD (left, blue) and SD (right, red) conditions for different gene sets with specific phases or maximum expression time points. Under LD conditions no significant difference is observed whereas under SD conditions genes with phases during the skotoperiod (dark period ZT8, ZT12, ZT16 and ZT20) present significantly longer offsets when compared to those genes with phases during the photoperiod (light period ZT0 and ZT4) according to Mann-Whitney-Wilcoxon test. **(F)** Boxplot illustrating how genes involved in different biological processes according to their gene ontology (GO) annotation present distinct protein/gene offsets that are longer under SD (red) than LD (blue) conditions. DNA replication (GO:0006260), photosynthesis (GO:0009521) and translation (GO:0006412) are chosen as examples exhibiting short and long protein/gene offsets. **(G)** Protein abundance and gene expression profiles under LD and SD conditions for *Sister Chromatid Cohesion 1* (left, *ostta03g03780*, *SMC1*), *Photosystem I Light Harvesting Complex 2* (center, *ostta03g04920*, *LHCA2*) and *Ribosomal Protein S1* (right, *ostta02g04680*, *RPS1*). This illustrates how genes involved in DNA replication or photosynthesis present short gene/protein offsets whereas genes involved in translation present long offsets.

Transcript/protein phase offsets did not show any correlation or dependence on common indexes or properties computed from protein sequences such as amino acid composition, charge or hydrophobicity (Supplementary Fig. 11D). Similarly, transcript phase under LD conditions did not affect transcript/protein offset lengths. Nonetheless, under SD conditions transcript/protein offsets were significantly longer for transcripts peaking during the night or skotoperiod (ZT8, ZT12, ZT16 and ZT20) when compared to those genes with transcript phases during the day or photoperiod (ZT0 and ZT4). However, we found that genes involved in different biological processes identified by specific Gene Ontology (GO) terms presented distinct short or long transcript/protein phase offsets, Supplementary Fig. 11E. Specifically, on the one hand, the most representative biological processes with short offsets were DNA replication (GO:0006260) and photosynthesis (GO:0009521) both under LD and SD conditions, (Figure 4F, Supplementary Fig. 11F). *Sister Chromatid Cohesion 1* (*ostta03g03780*, *SMC1*) and *Photosystem I Light Harvesting Complex 2* (*ostta03g04920*, *LHCA2*), illustrate short transcript/protein offsets (Figure 4G, H). On the other hand, translation (GO:0006412) was the most representative biological process exhibiting long transcript/protein offsets. *Ribosomal Protein S1* (*ostta02g04680*, *RPS1*) exemplifies genes with long transcript/protein offsets.

### Physiological rhythms are underpinned by an orchestration between proteome and transcriptome rhythmicity

Rhythmic patterns have been described for multiple physiological processes under diel cycles consisting of alternating light and dark periods in microalgae, including *Ostreococcus* ^25, 56, 63, 80^. These processes encompass cell cycle progression, primary and secondary metabolism. In this study, we performed physiological measurements of these processes and integrated them with our transcriptomic and proteomic data in order to elucidate the temporal orchestration at the different molecular levels that underlies physiological dynamics. To achieve this, we employed flow cell cytometry, pulse amplitude modulation (PAM), measured enzymatic activity, and determined carotenoid and starch content using the same samples utilized in our transcriptomic and proteomic analyses, ensuring a comprehensive integrative analysis.

### Cell cycle progression

Ostreococcus cell cycle follows the typical sequence of phases observed in binary fission: Gap 1 (G1) phase, DNA Synthesis (S) phase, Gap 2 and Mitotic (G2|M) phase^53, 55, 56^. To estimate the distribution of cells in each phase over diel cycles under LD and SD conditions, flow cell cytometry was employed to determine cell DNA content. Rhythmicity was observed in all cell cycle phases under both LD and SD conditions, with a higher significance level observed under LD conditions compared to SD conditions. This finding aligns with the previously observed decrease in synchronization in SD entrained cultures (Supplementary Table 7). A reduction of approximately 24% was observed in the number of cells entering the S phase under SD conditions compared to LD conditions. In SD entrained cultures, these cells remained in either the G1 or G2 phase (Supplementary Fig. 12A). Furthermore, significant backward shifts of approximately 4 hours were observed in the cell cycle phases under SD conditions compared to LD conditions (Figure 5A). These observations are consistent with the observed backward shifts in the time points when transcript and protein abundances reach their maximum levels in response to photoperiod shortening. Key genes involved in different cell cycle phases were identified in the Ostreococcus genome^54^, along with the corresponding time points when their transcripts and proteins abundances reach maximum levels under both LD and SD conditions (Supplementary Table 7). Integration and comparison using violin plots were conducted between transcript and protein abundance profiles and the profiles of the cell percentage in the S phase (Figure 5B), G1 phase, and G2|M phase (Supplementary Fig. 12B). Clear temporal offsets were observed between transcript and protein expression, with gene expression preceding protein abundance levels. Moreover, a shorter temporal offset was observed between protein levels and the cell percentage in each cell cycle phase.

**Figure 5.**
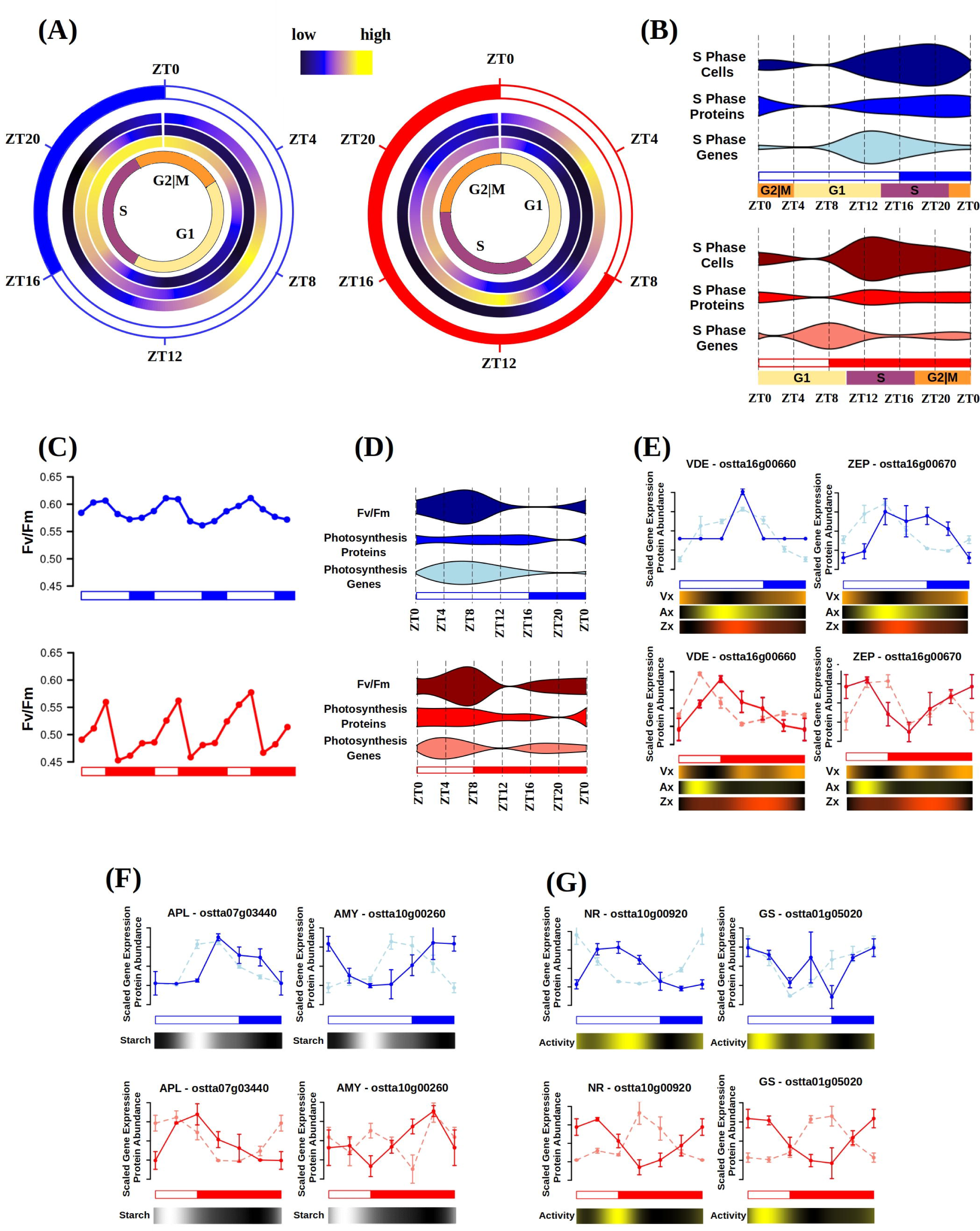
Rhythmicity of physiological measurements and integration with transcriptome and proteome dynamics under long and short day conditions. **(A)** Circular heatmaps representing the percentage of cells in each cell cycle phase over diel cycles. Black represents low percentage, blue medium values and yellow stands for high percentages. Left in blue, data under long day conditions (LD, 16h light / 8h dark), and right in red, data under short day conditions (SD, 16h light / 8h dark). ZTN, Zeitgeber time N, marks the time point N hours after dawn (lights on, ZT0). The outer circle represents photoperiods (light periods or days) in white, and skotoperiods (dark periods or nights) in blue or red for LD or SD conditions respectively. The next three circles represent the percentage of cells in the Gap 1 (G1), DNA synthesis (S) and Gap2 and Mitotic (G2|M) phases respectively. The innermost circle represents the different cell cycle phase temporal organization., G1 phase in yellow, S phase in purple and G2|M phase in orange. **(B)** Violin plots integrating S phase gene expression and protein abundance profiles with the percentage of cells in the S phase under LD conditions (top, blue) and SD conditions (bottom, red). Violin widths represent the corresponding average gene expression, protein abundance and percentage of cells in S phase profiles. White rectangles represent photoperiods (light periods or days), blue and red filled rectangles correspond to skotoperiods under LD and SD respectively (dark periods or nights). **(C)** Maximum quantum efficiency, Fv/Fm, profiles over three consecutive days under LD conditions (top, blue) and SD conditions (bottom, red) conditions. **(D)** Violin plots integrating Photosystem II transcript and protein abundance profiles with Fv/Fm data under LD conditions (top, blue) and SD conditions (bottom, red). **(E)** Transcript, light color, and protein, dark color, abundance profile for violaxanthin de-epoxidase (VDE, left) and zeaxanthin epoxidase (ZEP, right) under LD conditions (top, blue) and SD conditions (bottom, red). Heatmaps are incorporated below to represent the changes in violaxanthin (Vx, orange), antheraxanthin (Ax, yellow) and zeaxanthin (Zx, red) content. Black represents low content and the corresponding color, orange, yellow or red, represents high content. **(F)** Transcript, light color, and protein, dark color, abundance profile for ADP-glucose Pyrophosphorylase Large unit (APL, left) and Amylase (AMY, right) under LD conditions (top, blue) and SD conditions (bottom, red). Heatmaps are incorporated below to represent the changes in starch content. Black represents low content and white high content. **(G)** Transcript, light color, and protein, dark color, abundance profile for ADP-glucose Pyrophosphorylase Large unit (APL, left) and Amylase (AMY, right) under LD conditions (top, blue) and SD conditions (bottom, red). Heatmaps are incorporated below to represent the changes in starch content. Black represents low content and white high content.

Cyclins and Cyclins Dependent Kinases (CDKs) are essential components of the molecular machinery governing cell cycle progression, ensuring the proper timing and coordination of events in each phase^53^. Accordingly, we examined their gene expression patterns and protein abundance profiles, relating them to the temporal progression of cell cycle phases under LD and SD conditions (Supplementary Fig. 12 C, D and E). The gene expression level of *CyclinD* (*ostta18g01570*, *CYCD*) was found to increase from the beginning to the middle of the G1 phase (ZT4 – ZT9 under LD; ZT0 – ZT5 under SD) promoting cell cycle entry and progression to the S phase. During the second half of the G1 phase (ZT9 – ZT14 under LD; ZT5 – ZT10 under SD) and the first half of the S phase (ZT14 – ZT18 under LD and ZT10 – ZT14 under SD), *CyCD* expression decreased, while the gene expression of *Cyclin B* (*ostta01g06150, CYCB*) increased, inducing progression towards G2|M phase. A similar pattern was observed for both gene expression and protein abundance of *Cyclin Dependent Kinase B* (*ostta15g00670*, *CDKB*) under LD and SD conditions, with no apparent temporal offset between transcript and protein levels. *Cyclin A* (*ostta02g00150*, *CycA*) showed low gene expression rhythmicity, reaching its peak during G2|M phase at ZT0 under LD and ZT20 under SD conditions. Conversely, *Cyclin Dependent Kinase A* (*ostta15g00670*, *CDKA*) exhibited rhythmic gene expression and protein abundance under LD and SD conditions. In LD entrained cultures, *CDKA* gene expression peaked at the end of the G1 phase (ZT12), preceding the protein abundance peak at ZT20 during the S phase by 8 hours. However, in SD entrained cultures, *CDKA* gene expression and protein abundance peaked at ZT0 at the end of the G2|M phase with no apparent temporal offset. Genes involved in chloroplast division, such as *Filamentous Temperature-Sensitive Z* (*FtsZ*, *ostta07g01610*), play central roles in the G2|M phase showing gene expression peaks at the end of the G1 phase (ZT12 in LD and ZT8 in SD) preceding the protein peaks reached during the transition S/G2|M phase (ZT0 in LD and ZT16 in SD) (Supplementary Fig. 12F). Confocal microscopy images validated these findings by identifying cells with two chloroplasts as a result of recent divisions at ZT20 under LD conditions and ZT16 under SD conditions (Supplementary Fig. 12G).

### Photosynthesis and starch biosynthesis

The photosynthetic molecular machinery must dynamically adapt to seasonal variations in diel cycles to optimize light energy capture, utilization and ensure efficient carbon fixation and biomass production^81^. To assess the overall integrity and efficiency of Photosystem II (PSII), we measured the maximum quantum efficiency, Fv/Fm, using Pulse-Amplitude-Modulation (PAM) throughout diel cycles under LD and SD conditions (Figure 5C). Fv/Fm measurements exhibited rhythmicity under both LD (p-value 7.07× 10^-6^) and SD (p-value 2.02×10^-8^) conditions, peaking at ZT8 in both cases, albeit slightly lower (7%) under SD compared to LD conditions. In contrast, Fv/Fm reached its minimum at the end of the photoperiod (ZT16) under LD but at the beginning of the night (ZT12) under SD, with a 19% reduction. Violin plots were used to compare Fv/Fm measurements with transcript and protein abundance profiles of genes associated with PSII (Figure 5D). Under LD conditions, no temporal offsets were observed between the time points of maximum transcript and protein abundances and the highest Fv/Fm values. However, under SD conditions, short temporal offsets were identified between the profiles. Specifically, the early increase in gene expression at the beginning of the night, observed in, for example, genes encoding components of the Oxygen Evolving Complex *PSII subunits O*, *P* and *Q* (*PsbO*, *ostta14g00150*; *PsbP*, *ostta14g02630*; *PsbQ*, *ostta16g01620*) resulted in a corresponding increase in protein abundance and Fv/Fm values during the second half of the night. Similar expression patterns resulting in profiles with two peaks under SD conditions were found in genes such as *Protein Electron Transfer C* (*PetC*, *ostta01g06610*), *Ferredoxin* (*Fd*, *ostta17g00310*) and *ATPase delta subunit* (*ATPD*, *ostta07g01350*) coding for key components of Cytochrome b6f, Photosystem I and ATP Synthase, Supplementary Fig. 13. These genes constitute examples of the emergence of two peaks under SD conditions, one induced by the photoperiod maintained only under LL and the other one by the skotoperiod maintained only under DD. Genes coding for key components on the Calvin Cycle such as *Glyceraldehyde-3-phosphate dehydrogenase A* (*GAPDHA*, *ostta01g01560*) and *Fructose-1,6-bisphosphate aldolase* (*FBA*, *ostta01g03040*) are examples of *bona fide* circadian genes exhibiting rhythmicity under LD, SD, LL and DD. In all these examples no or short temporal offsets between transcript peaks (ZT8 under LD and ZT4 under SD) and protein peaks were found, Supplementary Fig. 13.

Glucose, a final product of the Calvin Cycle, can be stored long-term as starch. To further investigate the orchestration between transcript and protein abundances with physiological measurements, we assessed starch content throughout diel cycles under LD and SD conditions. Starch content profiles exhibited rhythmicity under both LD conditions (p-value 1.2× 10^-4^), reaching a peak of 41.2% of dry weight (DW) at ZT8; and under SD conditions (p-value 1.38×10^-4^), peaking at ZT4 with a value of 33.4% of DW (Supplementary Table 8). We observed a clear temporal sequence in gene activation preceding protein accumulation, with temporal offsets of approximately 4 hours for the enzymes involved in starch metabolism, Supplementary Fig. 13. Specifically, the first committed step in starch metabolism, the conversion of Glucose-1-phosphate into ADP-glucose, is catalyzed by ADP-glucose pyrophosphorylase, with its small and large subunit genes (*APS*, *ostta07g03070* and *APL*, *ostta07g03440*) being fully activated at ZT8 and ZT4 under LD and SD conditions, respectively. These activations preceded the peak of the corresponding proteins by approximately 4 hours. Subsequent steps involve the synthesis of amylose by Granule Bound Starch Synthase (*GBSS*, *ostta06g02940*) and amylopectin by Starch Branching Enzyme (*SBE*, *ostta03g00870*). The corresponding genes peaked at ZT12 under LD and presented two peaks at ZT20 and ZT4 under SD, while the encoded proteins reached their maximum abundances at ZT16 both under LD and SD conditions. The elongation of both amylose and amylopectin is performed by Soluble Starch Synthase (SSS, ostta16g02480) which showed its gene expression peak at ZT16 and ZT12, shortly preceding protein abundance, under LD and SD conditions, respectively. Starch homeostasis and degradation involve the enzymes *Debranching enzyme* or *Isoamylase* (*ISA*, *ostta02g05060*) and *Amylase* (*AMY*, *ostta10g00260*), which produce maltodextrin and maltose. The genes encoding for these proteins reached their maximum expression at ZT16 under LD and ZT8 under SD, preceding the accumulation of the corresponding proteins that reached their maximum during the night, Supplementary Fig. 13.

By integrating these gene expression and protein abundance patterns with starch content profiles over LD and SD diel cycles, we observed that starch accumulation is the result of a balance between synthesis and degradation. Under both LD and SD conditions, starch content reached its maximum at midday (ZT8 under LD and ZT4 under SD), despite the enzymes involved in starch biosynthesis and their corresponding genes peaking several hours later, toward end of the day. The halt in starch increase and the subsequent decrease in its content can be attributed to the activation of the genes encoding enzymes involved in starch catabolism or degradation, and the increase in abundance of the corresponding proteins during the second half of the day both under LD and SD conditions (Figure 5F).

### Carotenoid biosynthesis

Carotenoids are pigments that play a central role in photosynthetic organisms, including phytoplankton, as essential components of the molecular machinery involved in light harvesting and photoprotection^82^. To understand the adaptive nature of carotenoid content to seasonal variations in diel cycles and its implications for optimizing light energy capture and photoprotection, we examined the transcript and protein abundance profiles of carotenoid biosynthesis genes in Ostreococcus. These profiles were integrated with carotenoid content profiles, Supplementary Fig. 14A.

Carotenoid biosynthesis begins with the methylerythritol phosphate (MEP) pathway, responsible for producing the carotenoid precursors isopentenyl diphosphate (IPP) and dimethylallyl diphosphate (DMAPP). Most genes encoding enzymes involved in the MEP pathway, from the *1-deoxy-D-xylulose 5-phosphate synthase* (*DXS*, *ostta02g01730*) as the first committed step, to *4-hydroxy-3-methylbut-2-en-1-yl diphosphate reductase* (*HDR*, *ostta08g01180*) producing IPP and DMAPP peaked at ZT0 under LD and at ZT20 under SD, preceding the corresponding protein abundance peaks by up to 4 hours. Similar patterns were observed for the first enzymes in the carotenoid biosynthesis pathway leading to the committed step catalized by *Phytoene Synthase* (*PSY*, *ostta05g03530*).

Carotenoid biosynthesis can follow two different branches, alpha and beta, depending on the activity of *Lycopene epsilon/beta cyclase* (*LCYε/β*, *ostta14g00700*). The profiles of alpha-branch carotenoids, α-carotene, dihydrolutein, lutein, prasinoxanthin, micromonal and uriolide, as well as beta branch carotenoids, β-carotene, zeaxanthin, antheraxanthin, violaxanthin and neoxanthin, were determined over complete diel cycles under LD and SD conditions. All quantified carotenoids exhibited rhythmicity, except for violaxanthin, neoxanthin and uriolide under SD condtions (Supplementary Table 9).

The xanthophyll cycle, the interconversion between violaxanthin, antheraxanthin and zeaxanthin as a response to light intensity^83^, was especially active under LD conditions, although its activity was also detected under SD conditions. The changes in these xanthophylls coincided with the accumulation of transcripts and proteins encoded by genes associated with the xanthophyll cycle, with short temporal offsets (Figure 5E). Specifically, under LD conditions, violaxanthin decreased sharply during the first half of the day (ZT0 – ZT8), while antheraxanthin and zeaxanthin increased, corresponding to the accumulation of *Violaxanthin De-Epoxidase* (*VDE*, *ostta16g00660*) transcripts and proteins with a short temporal offset. Subsequently, during the second half of the day and night, antheraxanthin and zeaxanthin decreased, and violaxanthin increased, in agreement with the increament in gene expression and protein abundance of *Zeaxanthin Epoxidase* (*ZEP*, *ostta16g00670*).

A similar cycle was observed among the alpha carotenoids, with prasinoxanthin, the most abundant carotenoid in Ostreococcus, decreasing as ligh intensity increased, concomitant with an accumulation of both lutein and dihydrolutein. The enzymes involved in the interconversion of these carotenoids remain to be identified, and a comparison of their gene expression and protein accumulation was not feasible in this study.

### Nitrate assimilation

In all previously studied physiological processes short temporal offsets were identified between transcript and protein maximum abundances. In this section we examine the nitrate assimilation pathway in Ostreococcus, a physiological process that exhibited long temporal offsets between transcripts and proteins. Nitrate assimilation is a fundamental process in the metabolism of photosynthetic organisms, including microalgae, as it provides a key source of nitrogen for cellular growth and biomass production^84^. To study the adaptive response of nitrate assimilation to seasonal variations in diel cycles and its implications for optimizing nutrient uptake and metabolism, we examined the transcript and protein abundance profiles of this pathway in Ostreococcus (Supplementary Fig. 15).

Nitrate assimilation initiates with the uptake of nitrate by *Nitrate Transporters 2 and 3* (*NRT2*, *ostta10g00950* and *NRT3*, *ostta10g00940*), followed by its reduction to nitrite by *Nitrate Reductase* (*NR*, *ostta10g00920*). Nitrite is further reduced to ammonia by *Nitrite Reductase* (*NIR*, *ostta10g00930*). The central part of nitrate assimilation is played by the *Glutamine Synthetase* (*GS*, *ostta01g05020*) and *Glutamine Oxoglutarate Aminotransferase* (*GOGAT*, *ostta14g01900*) cycle, which converts inorganic nitrogen, ammonia, into glutamine and glutamate, a central precursor for the biosynthesis of nitrogen-containing compounds such as amino acids and nucleotides.

Under LD conditions, gene expressions of *NRT2/3*, *NR* and *NIR* reached their maximum at dawn (ZT0), while their protein abundances peaked 8 hours later at midday (ZT8), coinciding with the time point of maximum light irradiance. *GS* and *GOGAT* gene expressions and protein abundances peaked at the beginning of the day (ZT4) without noticeable temporal offsets. However, a slight increase in protein abundance was detected at the end of the day (ZT12) for both GS and GOGAT, Supplementary Fig. 15. In contrast, under SD conditions, all genes encoding the transporters and enzymes involved in nitrate assimilation showed their maximum expressions at midnight (ZT20), preceding their protein abundance peaks reached at dawn (ZT0), or midday (ZT4) by 4 or 8 hours. Notably, *GOGAT* gene expression displayed a bimodal pattern under SD conditions, maintaining the peak observed under LD at ZT4 besides the new peak at ZT20. Therefore, *GOGAT* gene expression constitutes an example of the emergence of complex expression patterns under SD conditions consisting of two gene expression peaks per day (Supplementary Fig. 15).

To validate these observed temporal offsets between transcripts and proteins, we measured the enzymatic activities of NR and GS throughout complete diel cycles under LD and SD conditions. Enzymatic activities profiles were significantly rhythmic (Supplementary Table 10). NR enzymatic activity reached its maximum at ZT8 under both LD and SD conditions coinciding with the protein abundance peak under LD and exhibiting a short forward temporal offset under SD. Similarly, GS enzymatic activity peaked at ZT0 under both LD and SD concomitant with the protein abundance peaks (Figure 5G).

## Discussion

Seasonality plays a key role in the natural geographical growth dynamics of marine phytoplankton, including the model microalgae Ostreococcus tauri^18^. Despite its expected impact in improving phytoplankton outdoors mass cultivation at industrial level, the molecular mechanisms underpinning the responses to seasonal variations in diel cycles remain to be characterized^10, 85^. In this study, we adopted a multiomics approach to unravel the adaptive orchestration between transcriptome and proteome rhythms governing cyclic physiological responses to seasonal changes in photoperiod length. Specifically, transcriptome and proteome dynamics were integrated with physiological measurements under summer long day (LD, 16h light: 8h dark) and winter short day (SD, 8h light: 16h dark) conditions.

Our transcriptomic analysis revealed that seasonality had no effect over the identity of rhythmic genes, with nearly identical cycling gene sets detected under LD and SD conditions, encompassing 80% of the transcriptome. This high transcriptome rhythmicity is consistent with previous findings under neutral day conditions (ND, 12h light: 12h dark) for Ostreococcus^51^ and other chlorophyte microalgae such as Chlamydomonas reinhardtii^72^. The non-rhythmic genes were significantly associated with stress responses and exhibited low expression or complete repression. However, these genes could potentially become rhythmic once highly expressed under the corresponding stress condition, resulting in a fully rhythmic transcriptome. Our result indicates that transcriptome rhythmicity in chlorophyte phytoplankton is much higher than the ones found in other organisms such as *Arabidopsis thaliana* 30-50%^28^, *Solanum tuberosum* 18-45%^86^, *Drosophila melanogaster* 24%^30^ or *Mus musculus* 3-10%^29^. Despite the high transcriptome rhythmicity under LD and SD diel cycles, only a small fraction (less than 5%) of the transcriptome comprised *bona fide* circadian genes that remained rhythmic even under free-running conditions of constant light (LL) and constant dark (DD) after LD and SD entrainment. These *bona fide* circadian genes were found significantly involved in photosynthesis and chloroplast organization, indicating that only these crucial processes are predominantly controlled by the autonomous circadian clock in Ostreococcus. This finding suggests that gene rhythmicity is strongly influenced by alternating light/dark cycles in this picoeukaryote. For instance, genes involved in essential processes, such as DNA replication, maintained rhythmicity under LL but were strongly repressed under DD, while genes involved in ribosome biogenesis exhibited rhythmicity under DD but lost it under LL. Generally, LL had a more detrimental effect on transcriptome rhythmicity (36.6% and 20% after LD and SD entrainment) than DD (43.2% and 52.2% after LD and SD entrainment), particularly in SD entrained cultures. These reductions in rhythmicity are consistent with LL free-running conditions studies in plants *Medicago truncatula*^31^, *Solanum tuberosum*^86^ and *Hordeum vulgare*^87^. However, free-running conditions consisting of constant dark (DD) remain to be explored at the transcriptomic level in plants. In contrast, in animals such as *Drosophila melanogaster* and *Mus musculus*, although having a low percentage of rhythmic genes, do not exhibit substantial reductions in rhythmicity under DD free-running conditions, indicating that most rhythmic genes are indeed circadian in these organisms^30, 88^.

Notably, rhythmic patterns under free-running conditions showed significant differences compared to those observed under LD and SD conditions. A reduction in amplitude was detected under LL, possibly due to decreased culture synchrony at the transcriptomics level. Individual cell transcriptional programs could be desynchronized by constant light, leading to largely out of phase individual gene expression profiles, resulting in damped average gene expression oscillations at the entire cell culture level. Similar desynchronization effects at the single cell level have been reported in Arabidopsis leaves under constant light^74, 75^, consistent with the autonomous character of circadian clocks. Conversely, a slight increase in amplitude was observed under DD conditions after SD entrainment, indicating a synchronization increase between individual cell transcriptional programs. This supports the notion that LL has a stronger desynchronization effect than DD free-running conditions in Ostreococcus cultures. Moreover, our analysis of the rhythmic patterns under free-running conditions unveiled typical responses of nocturnal character, such as backward and forward phase shifts under DD and LL, respectively, when compared to LD and SD. Similar responses have been observed in behavioral traits in nocturnal animals^76, 77^. Indeed, in both LD and SD entrained cultures, most transcripts peaked during the night or skotoperiod, confirming this nocturnal transcriptomic character.

Although the rhythmic genes were almost identical under LD and SD conditions, we observed significant seasonal effects on their dynamic profiles. Specifically, our data revealed backward phase shifts and reductions in amplitude under SD when compared to LD. Similar phase shifts in response to photoperiod shortening have been demonstrated at the level of individual genes in the model chlorophyte microalgae *Chlamydomonas reinhardtii*^89^. The reduction in amplitude could be attributed to an increased desynchronization of individual cells under SD when compared to LD conditions, as discussed earlier. These transcriptomic seasonal responses to changes in photoperiod length were captured in a predictive model, which allows us to estimate the phase and amplitude of any gene expression profile over the year. This model is based on a linear interpolation between the phases and amplitudes detected under SD and LD conditions. To validate this model, we used microarray data under ND conditions available for Ostreococcus. Based on our results, we concluded that approximately two thirds of the rhythmic genes with a single peak under both LD and SD conditions respond to seasonal changes in photoperiod length by gradually adjusting their phases in accordance with the increasing or decreasing length of the day over the year.

Another response to photoperiod shortening that we observed in our data was the emergence of rhythmic bimodal gene expression profiles with two peaks during the 24h cycle under SD conditions. The maintenance of only one of these peaks under LL and the other one under DD led us to hypothesize that the expression of these genes results from the combination of two distinct profiles, one controlled by the photoperiod and the other one by the skotoperiod. To model this response, we employed Non Linear Squares (NLS) to decompose bimodal expression profiles detected under SD conditions into two different profiles, and linear interpolation to simulate phase shifts of these two expression profiles as a response to changes in photoperiod. Similar to the previous finding, this model successfully predicted the emergence of gene profiles with two peaks in the microarray data generated under ND conditions for Ostreococcus as a validation. This model shows how under LD conditions the two distinct profiles regulated independently by the photoperiod and skotoperiod overlap in time producing a single peak while as the photoperiod shortens and the skotoperiod lengthens they become out of phase resulting in a bimodal profile under SD conditions. Bimodal rhythmic gene expression profiles has also been identified in plants^90^ and animals^91^ under specific conditions.

To further explore the seasonal responses in Ostreococcus tauri at the molecular level, we generated SWATH-MS proteomic data under LD and SD conditions and integrated them with the transcriptomic data described above. We successfully identified and quantified approximately 48% of the Ostreococcus annotated proteome under LD or SD conditions. While our proteome coverage over diel cycles was an improvement compared to previous proteomic studies in other model organisms, such as 12% in *Arabidopsis thaliana*^92^, 30% in *Drosophila melanogaster*^93^ and 9% in *Mus musculus*^94^, it was more modest compared to the recently reported 85% in *Ostreococcus tauri* under ND and LL conditions^68^. Nonetheless, the detected proteome was representative, covering most subcellular components or organelles. In contrast to the high transcriptome rhythmicity observed, only 55% of the proteome exhibited rhythmicity under LD or SD conditions, indicating a more dynamically cycling character in the transcriptome than in the proteome under diel cycles. In contrast to our results on transcriptome rhythmicity, seasonality had a major effect over the identity of the rhythmic proteins with only 9% of the detected proteome exhibiting rhythmicity independently of the photoperiod length. Due to the small fraction of the rhythmic proteome and the reported loss of rhythmicity under LL^68^, we decided not to explore SWATH proteomic data under LL and DD free-running conditions. Instead, we focused our analysis on integrating transcriptomic and proteomic dynamics over seasonal variations in diel cycles, specifically changes in photoperiod length.

Generally, we observed no coincidences between the corresponding rhythmic protein and transcript abundance profiles, indicating a lack of global correlation between proteome and transcriptome dynamics. Instead, we observed temporal offsets of several hours between the phases of protein and transcript abundance profiles. This observation pointed to the decoupling of transcription and translation and the existence of a significant regulation over translation initiation. However, when we aligned protein and transcript abundance profiles according to the detected temporal offsets, we found positive high correlation values, indicating that transcriptome and proteome dynamics are indeed linked but separated temporally. Similar temporal shifts between transcripts and proteins have been reported in other organisms, such as *Mus musculus*^95^ where over 50% of transcript phases were found to precede their corresponding protein phases by more than 6 hours.

In our study, we observed that temporal offset lengths were not uniform for all transcripts/proteins and showed no dependence on intrinsic protein properties that can be computed from amino acid sequences, such as charge or hydrophobicity. Instead, different temporal offset lengths were specifically associated with transcripts/proteins involved in particular biological processs. For instance, transcripts/proteins encoded by genes involved in photosynthesis exhibited short offsets, while long temporal offsets were observed for transcripts/proteins codified by genes related to translation. This finding indicates the existence of a differential posttranscriptional regulation under diel cycles for each specific biological process.

Notably, we also detected a seasonality effect on transcript/protein temporal offsets, with longer offsets observed under SD than LD conditions. Moreover, in SD entrained cultures, we found significantly longer temporal offsets for transcripts peaking during the skotoperiod compared to those reaching their maximum level during the photoperiod.

Finally, we integrated measurements of cyclic physiological responses to seasonal variations in diel cycles with the transcriptome and proteome dynamics discussed above, aiming to elucidate the full temporal orchestration at different molecular levels underlying the responses to changes in photoperiod length. Specifically, we investigated cell cycle progression, photosynthetic efficiency, starch accumulation, carotenoid content and enzymatic activities in nitrate assimilation. All these processes presented rhythmic patterns with backward shifts under SD compared to LD conditions, in agreement with similar transcriptome and proteome responses to photoperiod shortening.

In line with the described Ostreococcus algal blooms in spring and summer, in LD entrained cultures we found an increase in the number of cells entering S phase, enhanced photosynthetic efficiency as indicated by high Fv/Fm values, greater starch accumulations, and a more activated xanthophyll cycle compared to SD entrained cultures.

While we observed several hours of temporal offsets observed between transcripts and proteins, we found almost coincident protein abundance profiles and rhythmic physiological measurements of the corresponding biological process. These temporal offsets shed light on the physiological significance of the observed transcriptional programs over complete diel cycles. For instance, considering the temporal offset of 8 hours between the transcript and protein abundances of Nitrate Reductase, which also requires light for its activation. Under LD conditions, it is sufficient for the transcript to peak at dawn (ZT0) to make the time point of protein maximum abundance coincide with light maximum irradiance (ZT8). However, under SD conditions, with a photoperiod of only of 8 hours, to ensure that the protein peaks at ZT4 (the point of maximum light irradiance), the corresponding transcript must reach its maximum expression level during midnight (ZT20). This suggests the existence of molecular mechanisms that allow Ostreococcus tauri to adjust its transcriptome timing depending on the photoperiod length, considering the temporal offsets between transcripts and proteins to ensure that the proteins are available at the appropriate moment of the day. Such adaptive orchestration between transcriptome, proteome and physiological timing might play a crucial role in the ability of Ostreococcus tauri to thrive and optimize its physiological processes accordingly under seasonal variations of diel cycles.

## Methods

### Culture Conditions

*Ostreococcus tauri* sequenced strain RCC4221 was used for all experiments. The growth media was prepared using Artificial Sea Water (24.55g NaCl, 0.75g KCl, 4.07g MgCl_2_·6H_2_O, 1.47g CaCl_2_ ·2H_2_O, 6.04g MgSO_4_·7H_2_O and 0.21g NaHCO_3_ per 1L distilled water) supplemented with 1 mL of Solution I (100g NaNO_3_ in 250mL distilled water), 1mL of Solution II (700mg Na_2_HPO_4_ and 2.5g K_2_HPO_4_ in 250mL distilled water), 1mL of Solution III with trace metals (2.68g NH_4_Cl, 5.2g Fe-EDTA, 37.2g Na_2_-EDTA, 23mg ZnSO_4_, 14mg CoSO_4_, 7.89mg Na_2_ MoO_4_ ·2H_2_O, 2.5mg CuSO_4_, 1.7mg H_2_SeO_3_ and 180mg MnCl_2_·4H_2_O in 500mL distilled water) and 1mL of f/2 vitamin solution.

RCC4221 is routinely maintained at 20°C under LD conditions in flasks illuminated with white light providing approximately 50 μE m^-2^s^-1^ during the light period. When starting a new experiment one of such flasks was used to inoculate a roux culture bottle up to approximately 500 ml. This batch culture was grown at 20°C under continuous white light (50 μE m^-2^s^-1^) and continuously sparged with air supplemented with 1% CO_2_. After approximately two weeks, the culture from the roux bottle was used to inoculate two photochemostats. These consists of water jacketed bubble columns with 2 L capacity (7 cm diameter, 50 cm height) containing 1.8 L of cell suspension continuously sparged with air to ensure culture homogenization. The flow of water from the jacket to an external cooling device precisely controlled temperature at 20°C. A pH probe is submerged into the culture and connected to a pH meter serving as input to a LabJack that controls an electrovalve allowing the on demand injection of CO_2_ into the air stream entering the culture to maintain the pH at 8.0. Each photochemostat is illuminated during the corresponding light periods using six Phillips PL-32 W/840/4p white-light fluorescent lamps. Instead of abrupt dark-to-light and light-to-dark transitions our illuminating system controlled by a LabJack simulates the progressive increase and decrease observed in solar daylight cycles.

Each photochemostat is kept within a wooden case and covered by a completely opaque fabric to ensure that the illumination is only provided by our systems.

Initially after inoculation with the culture from the roux bottle the photochemostats were operated in batch mode for about 3-4 days with incident irradiance set to either LD or SD conditions depending on the experiment with maximal irradiance value fix to 1.5 mE m^-2^s^-1^. Subsequently, incident irradiance was progressively increased every day until reaching 2.5 mE m^-2^s^-1^. After approximately one week, the photochemostats were switched to continuous mode with a peristaltic pump adding fresh media continuously to the photochemostats during the light period at a flow rate of 45 mL h^-1^ during the photoperiod. Excess culture was removed at the same rate by the overflow in order to keep a constant volume. Once the culture reached stationary phase entrainment under LD or SD conditions was maintained up to four weeks before sample collection.

### Sample Collection, RNA Extraction and Purification

For each time point 50 mL of culture were collected for RNA extraction. Cells were pelleted by centrifugation (4 min, 5000 RCF, 4°C). The supernant was discarded and the pelleted cells were quickly resuspended in Phosphate-buffered saline solution and pelleted again by centrifugation (1 min, 5000 RCF, 4°C). After removal of the supernatant, pelleted cells were immediately flash frozen in liquid nitrogen and stored at −80°C.

Frozen pellets were resuspended in 400 μL of disruption buffer (García-Domínguez & Florencio, 1997) and directly added to a 1.5 mL Eppendorf tube (RNAse free and phenol-proof) containing 400 μL of phenol:chloroform 1:1 and 100 μL of acid washed glass beads (0.25–0.3 mm diameter; Braun, Melsungen, Germany). Mechanical disruption was performed by 30 min of repeated cycles of 60 s of vortexing and 60 s of incubating on ice. Extracts were centrifuged (4 °C) for 15 min at 13000 x g, producing three differentiated phases: an upper aqueous phase containing RNA, a white interphase containing DNA and a lower organic phase containing proteins, lipids and glass beads. The upper aqueous phase was collected and mixed with 400 μL of phenol:chloroform 1:1 and centrifugated in the same conditions only 5 min. This process was repeated three times more.In the last washed only chloroform was used to avoid phenol contamination of the RNA samples. The supernatant was incubated overnight at −20 °C in a solution of 80 μL 10 M LiCl and 550 μL 100% EtOH for RNA precipitationand finally, samples were centrifuged 10 min at 13000 x g 4 °C. Pellets were dried to avoid EtOH contamination.

Further RNA purification was performed using the Isolate II RNA Plant Kit (Bioline). Washing, DNase treatment and elution were carried out following the manufacturer instructions. The eluted RNA concentration and integrity were measured using a bioanalyzer 2100 (Agilent RNA 6000 Nano Kit).

### RNA-seq data generation and analysis

Library preparation was carried out following the manufacturer instructions and sequencing was performed on the Illumina NextSeq500 sequencer. Approximately 10 million 75nt long single end reads were generated for each sample. RNA-seq data was analysed using our pipeline MARACAS (MicroAlgae RnA-seq and Chip-seq AnalysiS) pipeline. Specifically, the high quality of the sequencing data was assessed using the software package FASTQC. The Ostreococcus tauri genome sequence and annotation v3.0 (https://mycocosm.jgi.doe.gov/Ostta4221_3/Ostta4221_3.home.html) were used as reference genome (Blanc-Mathieu et al., 2014). Reads were mapped to this reference genome with HISAT2 (Kim et al., 2019). Transcript assembly and gene expression estimation measured as FPKM (fragments per kilobase of exon and million of mapped reads) were performed using StringTie (Kovaka et al., 2019) and the bioconductor R package ballgown (Frazee et al., 2015). Principal components analysis and hierarchical clustering were performed using the R package FactoMineR (Lê et al., 2008).

### Sample Collection, Protein Extraction and Digestion

Sample collection was performed as described for RNA analysis. For cell disruption, 1 mL of Trizol, 100 μL of acid washed glass beads (0.25–0.3 mm diameter) and 40 μL of Protein Inhibitor Cocktail, PIC (25x) were added onto frozen pellets, followed by 3 disruption cycles (60 s agitation – 60 s incubation on ice) using a Mini-Beadbeater (BioSpec Products). Proteins were extracted using TRI Reagent (Sigma-Aldrich), according to the manufacturer’s instructions. The resulting proteins pellets were resuspended with 2mL of 0.3 M guanidine solution in 95% EtOH using 10 sonication cycles (30 s sonication - 30 s of incubating at 4 °C) and then centrifugated at 4 °C during 5 min at 8000 x g. This washing process was repeated twice, followed by two additional washing using 90% EtOH. The final pellets were resuspended in NH_4_HCO_3_ 50 mM/0.2% Rapidgest (Waters) and total proteins were quantified using a Qubit device. For each sample, 50 μg of proteins were incubated with dithiothreitol (DTT, final concentration 4.5 mM) for 30 min at 60 °C. Then, iodoacetamide to a final concentration of 10 mM was added and incubated for 30 min, under total darkness at room temperature. A treatment with trypsin was done overnight at 37 °C in a 1:40 trypsin:protein. After that, formic acid was added and incubated at 37°C for 1h. Finally, 2% acetonitrile (v/v) were added to reach a concentration of the digested sample around 0.5 μg of protein/μl of solution.

### SWATH-MS (sequential windowed acquisition of all theoretical mass spectra) proteomics data generation and analysis

SWATH-MS were performed on a time-of-flight TOF triple quadrupole hybrid mass spectrometer MS (5600 plus, Sciex) equipped with a nano electrospray source coupled to an nanoHPLC Eksigent model 425. The Sciex software Analyst TF 1.7 was used for equipment control and data acquisition. Peptides were first loaded onto a trap column (Acclaim PepMap 100 C18, 5 µm, 100 Å, 100 µm id × 20 mm, Thermo Fisher Scientific) under isocratical order in 0.1 % formic acid/2% acetonitrile (v/v) at a flow rate of 3 μL/min for 10 min. Subsequently, they were eluted on a reversed-phase analytical column, Acclaim PepMap 100 C18, 3 µm, 100 Å, 75 µm id × 250 mm, Thermo Fisher Scientific, coupled to a PicoTip emitter (F360-20-10-N-20_C12 from New Objective). Formic acid 0.1 % (v/v) was used as solvent A and 2% acetonitrile with formic acid 0.1 % (v/v) were used as solvent B. Peptides were eluted with a linear gradient of 5-35 % (v/v) of solvent B in 120 min at a flow rate of 300 nL/min. The source voltage was selected at 2600 V and the temperature was maintained at 100 °C. Gas 1 was selected at 20 PSI, gas 2 at zero, and curtain gas at 25 PSI.

For proteins identification, Data Dependent Acquisition DDA method were used. It consisted of a TOF-MS with a scan window of 400-1250 m/z (accumulation time of 250 ms) followed by 50 MS/MS with a scan window of 230-1500 m/z (accumulation time of 65 ms) and with a cycle time of 2574 s.

The spectral library was constructed by making one run with a mixture of the biological replicates corresponding to each time point (ZT0, ZT4, ZT8, ZT12, ZT16, ZT20) with the DDA method described. ProteinPilot v5.0.1 software (Sciex) was used to identify the proteins in the library. A pooled search of all runs was performed. The parameters of the Paragon method were: trypsin as enzyme and iodoacetamide as cysteine alkylating agent.

The Ostreococcus tauri annotated proteome v3.0 file from https://mycocosm.jgi.doe.gov/Ostta4221_3/Ostta4221_3.home.html linked to a Sciex Contaminants database were used in library construction. A false positive analysis (FDR) was performed and those with FDR < 0.01 were considered.

For each sample, the equivalent of 1 µg of digested protein was injected into each run. Before that, a standard (MS synthetic peptide calibration kit from Sciex) was injected to self-calibrate the equipment, control the sensitivity and chromatographic conditions. The described DDA method was used for SWATH runs with 60 ms of accumulation time and 3.7 s of cycle time. Three technical replicates for each one of the three biological replicates were analysed resulting in 9 replicates per time point.

The library generated by DDA (1% FDR) was used in the analysis using the Sciex software PeaKView 2.2 with the microapp SWATH 2.0, together with the data obtained from the SWATH runs. Using this program, the chromatographic traces of the ions were extracted and dumped into the Marker view 1.2.1.1 program where the list of identified proteins with their corresponding areas were generated. The parameters for extraction of ions and obtaining the areas were: 10 peptides per protein, 7 transitions of each peptide, threshold of confidence of the peptides set at 90 and FDR 1%. The software NormalyzerDE 1.6.0 (Willforss et al., 2019) was used to performed Quantile normalization. Data were imputed with mean imputation method between the nine technical and biological replicates.

### Identification of genes, proteins and physiological measurements exhibiting rhythmic patterns

The bioconductor R package RAIN (Rhythmicity Analysis Incorporating Non-parametric Methods) (Thaben and Westermark, 2014) was used to identify genes, proteins and physiological measurements exhibiting rhythmic expression patterns. A p-value threshold equal to 0.05 was used in all the cases under study. Simple rhythmic expression patterns with a single peak over a diurnal cycle were identified by setting the period parameter to 24 hours. More complex rhythmic expression patterns exhibiting two peaks over a diurnal cycle were identified by setting the period parameter to 12.

Significant differences between specific features not related to rhythmicity were assesed using the Mann-Whitney-Wilcoxon non parametric test implemented in the R function wilcox.test.

### Statistical comparison between different rhythmic patterns

Rhythmic patterns were fitted to a co-sinusoidal curve characterized by three parameters: mesor, amplitude and phase. The statistical significance of the differences in mesor, amplitude and phase between different groups was performed using the R package CircaCompare (Parsons et al., 2020). A p-value threshold of 0.05 was used to determine statistically differences. The significance of the global differences in these parameters was assessed using the Mann-Whitney-Wilcoxon non parametric test implemented in the R function wilcox.test.

### Functional Annotation of Gene Sets

Functional enrichment analysis over different gene sets was performed using our online tool AlgaeFUN (microALGAE FUNctional enrichment tool) that in turn is based on the bioconductor packages clusterProfiler, enrichplot and pathview; and the functional annotation package developed by our group org.Otauri.eg.db for *Ostreococcus tauri*.

### Cell cycle analysis: sample collection, cells fixation and staining, data acquisition and processing

A volume of 1.5 mL of cell suspension were harvested for each time point and diluted 1:10 in PBS. Two mL of these dilutions were centrifugated and cells in the pellets were fixed with 10 mL of 100% EtOH before storage at −20°C for, at least, 24h. After fixation, cell suspensions were centrifuged for 5 min at 3500 x g (room temperature) and resuspended in 1 mL of PBS, washed once with PBS and sonicated for 3 minutes in an Ultrasonic Cleaner (JSP, US21, ultrasonic power 50W), in order to eliminate cell clumps and aggregates before staining. In the staining process, 2μL of the Vibrant Dye Cycle Green (V35004, Thermo Fisher) (10 μM final stain concentration) were added to each sample and incubated 30 min (37°C) for selective DNA labeling. After incubation, cells were washed and transferred to flow cytometry tubes for cell cycle analysis. Flow Cytometry acquisition were performed with a BD FACS Canto II (BD Biosciences) where stained DNA were excited by a 488nm laser and emission was collected in a 530/30 nm PMT. Flow rate was low and linear amplification were established for the acquisition. Data were analyzed using FlowJo v.10.6.1 (Becton Dickinson & Company BD). Analysis was performed using one of the univariate cell cycle platform that FlowJo provides, specifically the Watson pragmatic algorithm (Watson et al., 1987) was used to adjust the data to the model.

### Photosynthetic activity: sample collection and data acquisition

Fresh culture was harvested at the different specific times of the day. The samples were diluted 1:1 with growing medium and incubated at 20 °C in total darkness during 10 min. In order to analyze photosynthetic parameters, Pulse-Amplitude-Modulation PAM fluorometry measurements were performed using a Waltz DUAL-PAM-100. After darkness incubation, the non-actinic modulated light (450 nm, 2.8 μE m^-2^ s^-1^) was turned on, in order to measure F_0_ (fluorescence basal level). Then, to determine F_m_ (the maximum fluorescence level), a saturating red light pulse of 655nm and 5000 μE m^-2^ s^-1^ was applied to the sample during 400 ms. Fv/Fm, that corresponds to the maximum potential quantum efficiency of Photosystem II when all reaction centers were open, was calculated as Fv/Fm = (F_m_ – F_0_)/F_m_.

### Starch content determination

At each time point, 50 mL of fresh culture was harvested and centrifuged at 7000 x g during 10 min. Then pellets were washed with 1% ammonium formiate (p/v) to eliminate salts from the growing medium and lyophilized. Approximately 2-3 mg of lyophilized biomass were added to hermetic tubes containing 1 mL of glass beads (0.25–0.3 mm diameter) and 2 mL of chloroform:methanol (2:1). Three disrupting cycles (60 s agitation - 60 s incubation on ice) were applied using Mini-Beadbeater (BioSpec Products). Then cellular extracts were separated from the beads and saved in new tubes. Cellular extracts were centrifuged for 4 min at 13000 x g and the supernatant was discarded. The addition of chloroform:methanol (2:1) and centrifugation steps were repeated until the pellets were white in order to ensure the elimination of pigments and lipids that could disturb the determination process. Finally, pigment free pellets were dried. Proposed protocol for plants in (Rufty & Huber, 1983) was adapted to Ostreococcus tauri.

Starch granules in dry pellets were alkaline solubilized with 1mL of 0.2 M KOH and heated at 100 °C. After 30 min, samples were gradually cooled and pH was adjusted to 5.0 adding 300 μL of 1 M acetic acid.

To starch digestion, 7.4 U of α-amylase were added and incubated 30 min at 37 °C, breaking down starch in small linear and branched oligosaccharides. After that, 5 U of amyloglucosidase were added and incubated 1-2 h at 55 °C releasing glucose residues. Finally, in order to stop enzymatic reaction, the samples were incubated at 100 °C 2 min and centrifuged at 13000 x g for 10 min discarding pellet. Enzymes were prepared in 0.1 M of sodium acetate pH 4.5. The quantification of released glucose residues from starch was achieved following (Rufty & Huber, 1983) protocol. The method combined two enzymatic activities: hexoquinase, that phosphorylated glucose residues, and glucose-6-phosphate dehydrogenase (G6PDH) that reduced NAD+ oxidizing the phosphorylated glucose. NADH generated could be measured at 340 nm and corelated to glucose concentration in a ratio 1:1. To achieve that measurement, quartz spectrophotometer cuvettes were used containing: 100 μL of the sample, 500 μL of hexoquinase buffer (100mM HEPES pH 7.7, 10 mM MgCl2, 0.04% BSA p/v, 1mM DTT), 100 μL of ATP mix (containing 10mM ATP diluted in Hepes 100 mM ph 7.7), 100 μL of NAD+ mix (containing 4 mM NAD+ diluted in Hepes 100 mM ph 7.7), 2 μL of 2.5 U μL-1 glucose-6-phosphate dehydrogenase and 200 μL of destile water. The absorbance of that mixture was measured at 340 nm followed by the addition of 5 μL of 1 U μL^-1^ hexoquinase enzyme and the incubation at 27 °C for 20 min. After that, a second measure allowed to determine the amount of NADH produced during the reaction. NADH absorbance was related to the amount of glucose residues and consequently to the initial amount of starch in each the sample from calibration curves with commercial starch.

### Carotenoid content determination

Four milligrams of lyophilized biomass were added to hermetic tube containing 1 mL of glass beads (0.25–0.3 mm diameter) and 1 mL of pure acetone. Three disrupting cycles (60 s agitation – 60 s incubation on ice) were applied using Mini-Beadbeater (BioSpec Products). Carotenoids extraction was achieved following (Del Campo et al., 2004) proposed method. Darkness was maintain during the entire process to avoid pigments degradation. After centrifugation 4 min at 13000 x g, cellular extract was collected and saved in new tube. Again, 1 mL of pure acetone was added to wash glass beads, centrifugated and collected the supernatant. This process was repeated until the supernatant turned colorless. Supernatants were joined in the same tube and acetone was evaporated using a stream of nitrogen gas. Finally, 350 μL of acetone were added for HPLC analysis. A Hitachi HPLC (Elite LaChrom), equipped with a photodiode-array detector (Hitachi L-2455) was used. Separation was performed on a Waters NovaPak C-18 (3.9×150 mm, 4 µm particle size, 60 Å pore size) column. Following proposed method, the eluents used to create a gradient through the mobile phase were: eluent A (0.1 M ammonium acetate and 15:85 v/v H2O-methanol) and eluent B (44:43:13 v/v methanol-acetonitrile-acetone). Temperature was maintained constant (20 °C) during the whole process and eluents flowed at 800 μL min^-1^. Different carotenoids were identified following retention times and absorption profiles of previous known carotenoids analyzed. Quantification was calculated as a percentage of the total peak area.

### Data and Code availability

RNA-seq data generated in this study is freely available from the Gene Expression Omnibus (GEO) database under the accession number GSE155535. The data analysis code developed using the statistical programming language R is freely available from the following GitHub repository: https://github.com/fran-romero-campero/SANDAL.

## Acknowledgements

This work was supported by BIO2017-84066-R (MINOTAUR) and PID2021-123984OB-I00 (ELECTRA) from the Spanish Ministry of Science and Innovation. We would like to acknowledge Eloisa Andújar and Mónica Pérez from the CABIMER Genomics Unit for their assistance with high-throughput sequencing, Rocío Rodríguez from the IBVF Proteomics Units for guidance with SWATH-MS proteomics, Carlos Parejo from the IBVF Chromatography and Mass Spectrometry Unit for his contribution in carotenoid content determination, Alicia Orea from the IBVF Microscopy Service for her help with microscopy imaging, Enrique Frías from the Cic-Cartuja Cell Culture Unit for his assistance with *Ostreococcus tauri* cultivation and José Moreno Fernández for initially setting up *Ostreococcus tauri* cultivation in photochemostats for this project.

## Author Contributions

ABRL, MEGG, MMP and MGG performed wet lab experiments. ABRL, CA and FJRC generated and analysed RNA-seq transcriptomic and SWATH-MS proteomic data. ABRL, MJCP and FJRC generated and analyzed cell cycle data. ABRL and FJRC generated and analysed photosynthetic efficiency measurements. ABRL, MEGG, MMP and CA measured and analyzed starch accumulation, carotenoid content and enzymatic activities. ABRL, MGG and FJRC interpreted the results and wrote the manuscript. All authors read and approved the final manuscript.

## Competing Interests statement

The authors declare that they have no competing interests.

## Notes

### Competing Interest Statement

The authors have declared no competing interest.

