## Supplemental Table 1 for "Multiomics responses to seasonal variations in diel cycles in the marine phytoplanktonic picoeukaryote *Ostreococcus tauri*"

| **Sample Name** | **Number of reads** | **Percentage of mapped reads to the nuclear genome** |
| --- | --- | --- |
| *LD ZT0 1* | 9,446,680 | 94.09% |
| *LD ZT4 1* | 9,123,089 | 84.33% |
| *LD ZT8 1* | 9,382,983 | 88.28% |
| *LD ZT12 1* | 11,090,892 | 93.81% |
| *LD ZT16 1* | 10,365,158 | 91.13% |
| *LD ZT20 1* | 9,611,641 | 94.90% |
| *LD ZT0 2* | 9,911,801 | 95.07% |
| *LD ZT4 2* | 10,373,968 | 92.62% |
| *LD ZT8 2* | 9,361,305 | 94.61% |
| *LD ZT12 2* | 11,531,047 | 96.77% |
| *LD ZT16 2* | 10,096,408 | 91.95% |
| *LD ZT20 2* | 9,840,739 | 95.96% |
| *LD ZT0 3* | 9,271,993 | 74.71% |
| *LD ZT4 3* | 12,219,220 | 73.30% |
| *LD ZT8 3* | 10,295,962 | 57.05% |
| *LD ZT12 3* | 10,776,249 | 73.88% |
| *LD ZT16 3* | 9,694,347 | 71.37% |
| *LD ZT20 3* | 9,150,451 | 86.44% |
| *LD ZT0 4* | 10,040,606 | 80.48% |
| *LD ZT4 4* | 13,076,912 | 98.51% |
| *LD ZT8 4* | 14,287,019 | 98.70% |
| *LD ZT12 4* | 14,578,626 | 98.48% |
| *LD ZT16 4* | 12,754,119 | 98.09% |
| *LD ZT20 4* | 11,247,222 | 92.83% |
| *LD ZT0 5* | 10,652,932 | 97.36% |
| *LD ZT4 5* | 10,829,705 | 97.46% |
| *LD ZT8 5* | 7,714,090 | 98.19% |
| *LD ZT12 5* | 12,194,496 | 98.34% |
| *LD ZT16 5* | 10,793,769 | 98.17% |
| *LD ZT20 5* | 12,165,205 | 97.94% |
| *SD ZT0 1* | 12,493,583 | 98.05% |
| *SD ZT4 1* | 13,622,449 | 98.27% |
| *SD ZT8 1* | 11,074,612 | 98.07% |
| *SD ZT12 1* | 11,907,980 | 96.64% |
| *SD ZT16 1* | 10,879,032 | 97.80% |
| *SD ZT20 1* | 11,149,175 | 97.88% |
| *SD ZT0 2* | 14,740,881 | 95.05% |
| *SD ZT4 2* | 12,296,826 | 97.94% |
| *SD ZT8 2* | 12,311,981 | 97.84% |
| *SD ZT12 2* | 11,499,122 | 96.45% |
| *SD ZT16 2* | 10,968,505 | 97.49% |
| *SD ZT20 2* | 12,299,619 | 97.76% |
| *SD ZT0 3* | 7,072,975 | 95.81% |
| *SD ZT4 3* | 11,914,969 | 84.17% |
| *SD ZT8 3* | 8,560,034 | 95.06% |
| *SD ZT12 3* | 8,898,970 | 97.89% |
| *SD ZT16 3* | 9,814,400 | 97.30% |
| *SD ZT20 3* | 9,370,821 | 95.88% |
| *SD ZT0 4* | 8,980,904 | 96.01% |
| *SD ZT4 4* | 9,370,821 | 95.88% |
| *SD ZT8 4* | 14,529,619 | 96.44% |
| *SD ZT12 4* | 11,373,712 | 98.13% |
| *SD ZT16 4* | 13,933,214 | 92.04% |
| *SD ZT20 4* | 9,836,745 | 97.33% |
| *SD ZT0 5* | 10,287,353 | 94.11% |
| *SD ZT4 5* | 9,758,563 | 97.61% |
| *SD ZT8 5* | 9,652,648 | 88.35% |
| *SD ZT12 5* | 10,409,149 | 87.66% |
| *SD ZT16 5* | 9,221,180 | 97.26% |
| *SD ZT20 5* | 8,460,013 | 93.39% |
